# Substrate Stiffness Reshapes Layer Architecture and Biophysical Features of Human Induced Pluripotent Stem Cells to Modulate their Differentiation Potential

**DOI:** 10.1101/2024.02.01.578393

**Authors:** Jack Llewellyn, Anne Charrier, Emmanuèle Helfer, Rosanna Dono

**Affiliations:** Aix Marseille Univ, CNRS, IBDM, Turing Centre for Living Systems, NeuroMarseille, Marseille France; Aix Marseille Univ, CNRS, CINAM, Turing Centre for Living Systems, 13009 Marseille, France

**Author notes:** Corresponding author **Correspondence to:** Rosanna Dono, Ph.D., Aix Marseille Univ, CNRS, IBDM, Turing Centre for Living Systems, 163 Avenue de Luminy, case 907 - 13009 Marseille (France).

**Keywords:** Human induced pluripotent stem cells (hPSCs), hPSC self-renewal and differentiation, Mesendoderm and Endoderm Differentiation, Silicone, Hydrogels, Substrate stiffness, Epithelial organization, Mechanoenvironment

## Abstract

Lineage-specific differentiation of human induced pluripotent stem cells (hiPSCs) relies on complex interactions between biochemical and physical cues. Here we investigated the ability of hiPSCs to undergo lineage commitment in response to inductive signals and assessed how this competence is modulated by substrate stiffness. We showed that Activin A-induced hiPSC differentiation into mesendoderm and its derivative, definitive endoderm is enhanced on gel-based substrates softer than glass. This correlated with changes in tight junction formation and extensive cytoskeletal remodeling. Further, live imaging and *in silico* studies suggested changes in cell motility, shape, forces and pressures, underlie hiPSC layer reshaping on soft substrates. Finally, we repurposed an ultra-soft silicone gel, which may provide a suitable substrate for culturing hiPSCs at physiological stiffnesses. Our results provide mechanistic insight into how epithelial mechanics dictate the hiPSC response to chemical signals, and provide a tool for their efficient differentiation in emerging stem cell therapies.

## Introduction

Human pluripotent stem cells (hPSCs), which include human embryonic stem cells (hESCs) and human induced pluripotent stem cells (hiPSCs), are characterized by their ability to self-renew indefinitely in culture while maintaining their ability to differentiate into all human cell types. These properties make them a valuable model system for studies on developmental biology and the genetic basis of diseases as well as for applications in fields such as pharmacology, tissue engineering, and regenerative medicine^1, 2^. Much research has been dedicated to exploring methods to induce the differentiation of these cells towards specific lineages by taking advantage of insights gained from developmental biology studies^3–5^.

In embryos, fate and differentiation of PSCs is controlled by a complex interplay of biochemical cues including growth factors, inhibitors, and small bioactive molecules present in their microenvironment^6–10^. The influence of these biochemical cues has been extensively studied and standard differentiation protocols *in vitro* mimic known stages in development by the timed addition of biochemical players. Many experimental studies and computational models have also revealed that hPSCs sense and respond to various physical cues encoded by the microenvironment, such as local forces, cell-cell contacts and matrix stiffness. With the development of engineering substrates and apparatus, the effects of these mechanical stimuli sensed by PSCs in normal developmental processes are currently being tested in order to provide the best approach for the efficient generation of differentiated cells with native properties.

Matrix stiffness, usually characterized by the Young’s modulus (one of several different elastic moduli), has emerged as an important physical factor regulating multiple aspects of PSC behavior including division, migration and differentiation. For example, studies on hPSC lineage entry have shown that soft substrates in combination with high mechanical tension enhance hESC mesodermal differentiation^11, 12^. Yet, substrates softer than tissue culture polystyrene plates (GPa range) induce hESCs and hiPSCs to exit the pluripotency stage and promote spontaneous up regulation of endodermal genes^13, 14^. In another study, soft substrates were shown to favor better maintenance of hESC-derived endodermal cells suggesting that control of substrate stiffness might enable a superior control of live cells’ endodermal properties^15^. Matrix stiffness, which matches the stiffness of native tissues, can also guide stem cell differentiation down corresponding tissue lineages. For instance, substrates with Young’s moduli close to those of brain (0.1 to 0.7 kPa), pancreas (1.4 to 4.4 kPa), muscle (8 to 17 kPa) and bone tissue (25 to 40 kPa) direct hPSCs to differentiate into neurons, beta cells, myotubes and osteoblasts, respectively^16–19^. Taken together, these studies suggest that substrate stiffness can provide a tunable parameter to promote PSC lineage entry and differentiation. However, how substrate stiffness interacts with other differentiation-inducing factors, such as biochemical factors, remains largely unclear.

With gradual uncovering of the effect of substrate stiffness on directing cell biological properties, the question of how cells sense their physical microenvironment and make adjustments to control cell function has also raised rapid interest. An emerging concept is that the matrix stiffness is transduced in distinct cell physical features such as cell fluidity, forces and cytoskeleton organization that may be essential for cellular identity and function^20^. For example, studies on chondrocytes have revealed that soft matrix can induce a reorganization of the actin cytoskeleton important enough to efficiently promote chondrocyte differentiation from ATDC5 chondroprogenitor cells^21^. Yet, the elasticity of differentiated chondrocytes increases with matrix stiffness in order to create a homeostatic balance^22^. Similarly to chondrocytes, cancer cells also adjust their viscoelasticity in response to changes in extracellular matrix stiffness in order to maintain their invasive phenotypes^23^. Although these and other findings highlight the importance of the biophysical interplay between cells and their microenvironment for cell identity and behavior, these feedback mechanisms remain largely unexplored.

In this study we examined the ability of hiPSCs to acquire a mesendodermal fate in response to mesendoderm-inducing signals and assessed whether tuning the substrate stiffness can enhance this competence. Our rationale was that modeling this early step of embryonic development *in vitro* could provide the best approach to produce mesendodermal-derived cell types with native properties. We also examined the matrix stiffness-dependent mechanosensing of hiPSCs with a focus on cell’s physical properties with the aim of providing a quantitative understanding of the mechanisms that regulate cell interactions with substrates of different stiffness. We showed that hiPSC lineage entry into mesendoderm and its derivative definitive endoderm in response to differentiation triggering factors is enhanced when undifferentiated cells are exposed to gel-based substrates with stiffnesses lower than that of standard glass supports. This correlated with changes in the dynamics of tight junction (TJ) formation and with an extensive remodeling of the actin cytoskeleton. Moreover, live imaging of cells as well as *in silico* inference studies revealed that the biophysical mechanisms underlying the enhanced differentiation potential of these hiPSCs exposed to softer substrates may rely on distinct cell motility, shape, forces and pressure. Given that some hiPSC lines do not grow on ultra-soft polyacrylamide (PA) gels, our results also provide evidence that silicone matrices called QGel can provide a suitable alternative for expansion and differentiation of hiPSCs on ultra-soft substrates. Our studies, by providing new knowledge of matrix stiffness-dependent biological and biophysical behavior of hiPSCs, have important implications for optimization of matrix material and for advancing hiPSC-based clinical applications.

## Results

### Efficiency of hiPSC lineage entry is cell matrix stiffness dependent

To evaluate whether matrix stiffness regulates hiPSC self-renewal and mesendodermal cell fate, we cultured hiPSCs on substrates of different stiffness. We fabricated substrates made of thin (50 to 100 µm thick), flat silicone gels (polydimethylsiloxane, PDMS) deposited on glass coverslips via high-speed spin coating. The Young’s modulus of these PDMS gels can be tuned by varying the base:catalyst (PDMS:crosslinker) ratio in the silicone mixture. The two ratios used here led to Young’s moduli of 2.18 MPa and 77.6 kPa, much lower than that of glass, which is in the GPa range (Table 1). Both PDMS gel layers and glass supports were coated with Matrigel to enable hiPSC attachment. We first examined whether changes in matrix stiffness affect hiPSC self-renewal by analyzing the expression of pluripotency markers. Immunocytochemistry analysis of SOX2, OCT4, and NANOG proteins revealed that all cells remained positive for and maintained high expression levels of these pluripotency genes regardless of matrix stiffness (glass, 2.18 MPa, and PDMS 77.6 kPa; Figure 1A). Analysis of OCT4 and of the epithelial marker E-Cadherin (E- CAD) showed unchanged expression levels across a large range of matrix stiffnesses going from that of glass (of the order of the GPa) to that of the less reticulated PDMS gel (49:1 base:catalyst, 77.6 kPa), suggesting that changes in substrate stiffness do not perturb expression of pluripotency genes (Figure S1A). Interestingly, quantitative analysis of cell confluency over time suggests an increase in cell density for cultures grown on PDMS gels compared to glass (Figures S1B and S1C; see Material and Methods).

**Table 1.** Gel composition and corresponding Young’s modulus with standard deviations. N: Number of measurements.

| Gel type/Young's modulus | Standard deviation | N | Composition |  |
| --- | --- | --- | --- | --- |
| PDMS |  |  | Base % | Cross-linker % |
| 2.18 MPa | 0.213 MPa | 33 | 90 | 10 |
| 77.6 kPa | 13.3 kPa | 36 | 98 | 2 |
| PA |  |  | Acrylamide % | Bis-acrylamide % |
| 32.1 kPa | 9.47 kPa | 36 | 12 | 0.2824 |
| 25.1 kPa | 5.08 kPa | 36 | 10 | 0.282 |
| 11.6 kPa | 1.91 kPa | 51 | 7.464 | 0.16 |
| 4.07 kPa | 1.42 kPa | 44 | 5.488 | 0.05 |
| 2.15 kPa | 0.512 kPa | 35 | 3 | 0.06 |
| QGel |  |  | Solution A % | Solution B % |
| 31.2 kPa | 2.5 kPa | 23 | 33 | 67 |
| 1.9 kPa | 0.19 kPa | 24 | 40 | 60 |
| <1.9 kPa | - | - | 56 | 44 |

**Figure 1.**
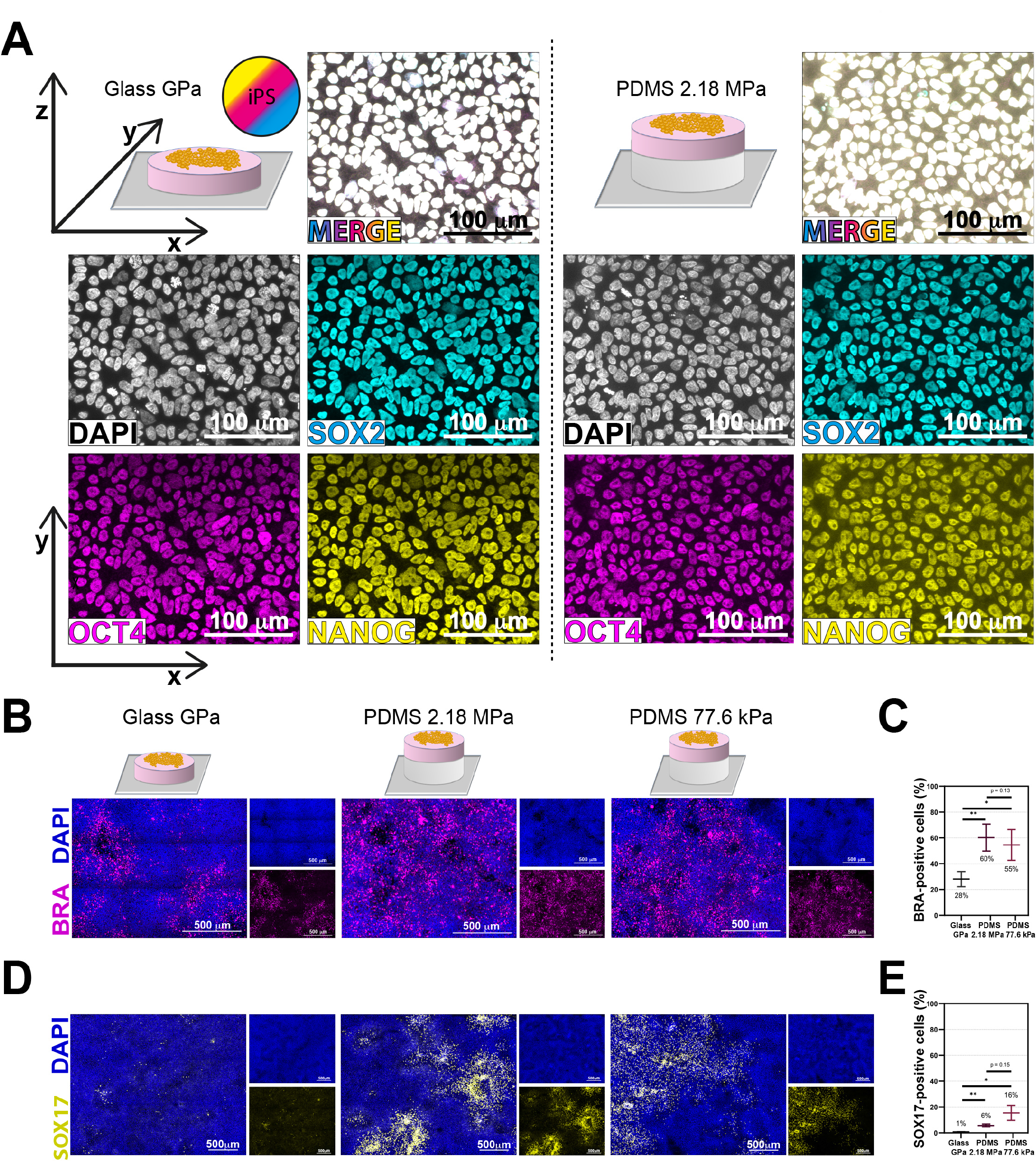
HiPSCs cultured on functionalized PDMS gel substrates preserve pluripotency while improving the efficiency of early lineage commitment. **(A)** Representative immunofluorescence images (IF) of the nuclear marker DAPI and of the pluripotency markers SOX2, OCT4, and NANOG in hiPSCs grown on glass (left) and on PDMS gel of stiffness 2.18 MPa (right). Cells were fixed and images were taken 24 hours after the removal of ROCK inhibitor (48 h after cell seeding at 110,000 cells/cm^2^). **(B)** Representative IF images of DAPI and of the mesendoderm marker T-Brachyury (BRA) in hiPSCs grown on glass (left), 2.18 MPa PDMS gel (middle), and 77.6 kPa PDMS gel (right). Cells were fixed and images were taken 24 hours after the addition of 100 ng/ml Activin A (48 hours after the removal of ROCK inhibitor, and 72 hours after cell seeding at 110,000 cells/cm^2^). **(C)** Quantification of the number of BRA-positive cells in the three stiffness conditions at the time point illustrated in **(D)**. BRA-positive cells were defined as those whose mean fluorescence intensity was above the 95^th^ percentile of the mean BRA intensity in Activin A-negative control cells at the same time point and stiffness. At least 10 randomly selected frames of view were taken per condition, covering >2.23 mm² in total. N = 7 paired replicates. **(D)** Representative IF images of DAPI and of the definitive endoderm marker SOX17 in hiPSCs grown on glass (left), 2.18 MPa PDMS gel (middle), and 77.6 kPa PDMS gel (right). Cells were fixed and images were taken 24 hours after the addition of 0.2% fetal bovine serum (FBS; 48 hours after the addition of 100 ng/ml Activin A, 72 hours after the removal of ROCK inhibitor, and 96 hours after cell seeding at 50,000 cells/cm^2^). **(E)** Quantification of the number of SOX17-positive cells in each of the three stiffness conditions at the time point illustrated in (**F**). SOX17-positive cells were defined via automatic thresholding. At least 3 randomly selected frames of view were taken per condition, covering >21.2 mm² in total. N = 4 unpaired replicates. Scale bar in images = (A) 100 µm and (B and D) 500 µm. Data are represented as mean ± SEM, and p values show the results of a two-tailed t test (*, < 0.05; **, < 0.01; ***, < 0.001).

Next, we tested the response of hiPSCs to the signal Activin A, an *in vitro* surrogate of Nodal, commonly used to trigger mesendoderm differentiation^24–28^. 24 hours following the addition of 100 ng/mL Activin A to the culture medium, we quantified the competence of cells to acquire a mesendodermal fate when exposed to different matrix stiffnesses by analyzing the mesendoderm marker T-Brachyury (BRA). Immunocytochemistry revealed that the number of BRA-positive cells increased by 2-fold in cells grown on PDMS gels compared to glass (Figures 1B and 1C; percentage of BRA-positive cells: glass 28.1%±5.8%; PDMS 2.18 MPa: 60.3%±10.4%; PDMS 77.6 kPa: 54.5%±11.9%). Interestingly, while significantly more BRA-expressing cells were seen when comparing either PDMS gel versus glass, no significant difference was seen between the two PDMS gels.
**

To investigate whether this increased differentiation efficiency in hiPSCs cultured on matrices softer than glass is maintained as mesendodermal cells acquire an endodermal fate, we applied a minimal endoderm differentiation protocol by culturing mesendodermal cells only in the presence of Activin A (instead of Activin A and the GSK3 inhibitor CHIR99021^29^) and of defined heat-inactivated 0.2% fetal bovine serum (FBS) for an additional 24 hours, and determined the extent of definitive endoderm by using the marker SOX17. Of note, a significant increase in the number of SOX17-positive cells was observed in the differentiating cells cultured on both PDMS gels compared to glass, which we quantified as at least a 6-fold increase in SOX17-positive cells (Figures 1D and 1E; percentage of SOX17-positive cells: glass 0.8%±0.1%; PDMS 2.18 MPa: 5.7%±1.2%; PDMS 77.6 kPa: 15.5%±5.8%). This quantitative analysis also revealed a tendency for cells grown on the 77.6 kPa PDMS gel to differentiate more efficiently into endodermal cells than those grown on the 2.18 MPa PDMS gel, thus suggesting a positive correlation between softer substrates and competence to become endodermal cells (Figures 1D and 1E). Our inability to accurately measure the difference between the number of SOX17-positive cells in hiPCs differentiated on glass versus those differentiated on gel may be due to the degree of variance we observed in the softest gel, as gels become less homogenous as the catalyst concentration is reduced to its lower limit^16, 30^.

The PDMS-silicone base utilized in the experiments described above (Figure 1) is well-documented and commonly used for cell culture applications, however a consistently reported lower limit of the base:catalyst ratio limits its use to study cell differentiation efficiency at very low stiffnesses <1 kPa^16, 30, 31^. In previous studies of mammalian cell behavior on ultra-soft substrates, polyacrylamide (PA) hydrogels have been commonly used. We therefore investigated the differentiation of hiPSCs on PA hydrogel substrates. Similar to PDMS gels, PA mixtures of acrylamide and bis-acrylamide were prepared at varying ratios to fabricate PA hydrogel of varying stiffnesses, with Young’s moduli of 2.15, 4.07, 11.60, 25.10, and 32.10 kPa (Table 1). In agreement with previous studies of hiPSCs/hESCs^32–34^, we found that our hiPSCs did not grow as a 2D epithelium at PA stiffnesses below 10 kPa and preferentially formed 3D aggregates (Figure S1D) even in the presence of increasing concentration of Matrigel or by gradually decreasing matrix stiffness to favor adaptation of cells to the new environment. Therefore, we concluded that the use of PA hydrogels was not compatible with studies of matrix stiffness effects on differentiation of our hiPSC line for Young’s modulus values below 10 kPa. However, we reasoned that comparing the mesendoderm differentiation properties of hiPSCs grown on PDMS gels versus PA hydrogels of comparable stiffness could provide insights into the effects of scaffolds with distinct surface chemistry. Our results highlighted that hiPSCs grown on PDMS and PA hydrogel of a few tens of kPa stiffness underwent both mesendoderm differentiation as assessed by the presence of BRA-positive cells, respectively (Figure S1E). Moreover, we did not observe any difference in differentiation efficiency thus showing that scaffold mechanics, not surface chemistry, is the main determinant of this differentiation efficiency (Figure S1F; percentage of BRA-positive cells: PA 11.6 kPa: 52%±6%; PDMS 77.6 kPa: 53%±6%).

Taken together these results show that substrates with a lower stiffness than glass are compatible with expansion of hiPSCs and efficient differentiation into mesendoderm. As the 2-fold increase of mesendoderm-positive cells translates into a 6-fold increase into SOX17-positive cells as differentiation proceeds (see the number of SOX17-positive cells in hiPSCs differentiated on PDMS gels), it is likely that an exposure of hiPSCs to soft matrices at the lineage entry point amplifies the capability of differentiating cells to pursue the later differentiation steps.

### Epithelial and cytoskeletal organization is modulated by substrate stiffness

Previously, we have shown that the strength of the hiPSC response to Activin A and the differentiation efficiency into the mesendoderm lineage can be enhanced by disrupting hiPSC epithelial integrity^35^. To understand whether changes in matrix stiffness alter the hiPSC epithelial layer, we used the TJ protein Zona Occludens 1 (ZO1) to map cell-cell contacts. 24 hours after the removal of Rho kinase inhibitor (ROCK inhibitor) – the time point at which Activin A was introduced in differentiation assays – cells exhibited a reasonable organized epithelial structure on the glass substrate. In contrast we found a significant higher loss of epithelial integrity in hiPSCs grown on both PDMS gels, whether of 2.18 MPa or 77.6 kPa stiffnesses (Figure 2A). Moreover, quantitative analysis revealed similar levels of disorganization independently of the PDMS gel stiffness (Figure 2B; percentage of ZO1 organization: glass 67%±7%; PDMS 2.18 MPa: 52%±8%; PDMS 77.6 kPa: 49%±6%). Of note, analysis of epithelial integrity by ZO1 immunostaining at later time points of 48 and 72 hours after ROCK inhibitor removal revealed a progressive reorganization of the epithelial cell layer in hiPSCs cultured on PDMS gels with an establishment of TJs to nearly 100% after 72 hours (Figure S2A). This result indicates that reducing matrix stiffness causes a delay in TJ formation rather than preventing this cell-cell interaction.

**Figure 2.**
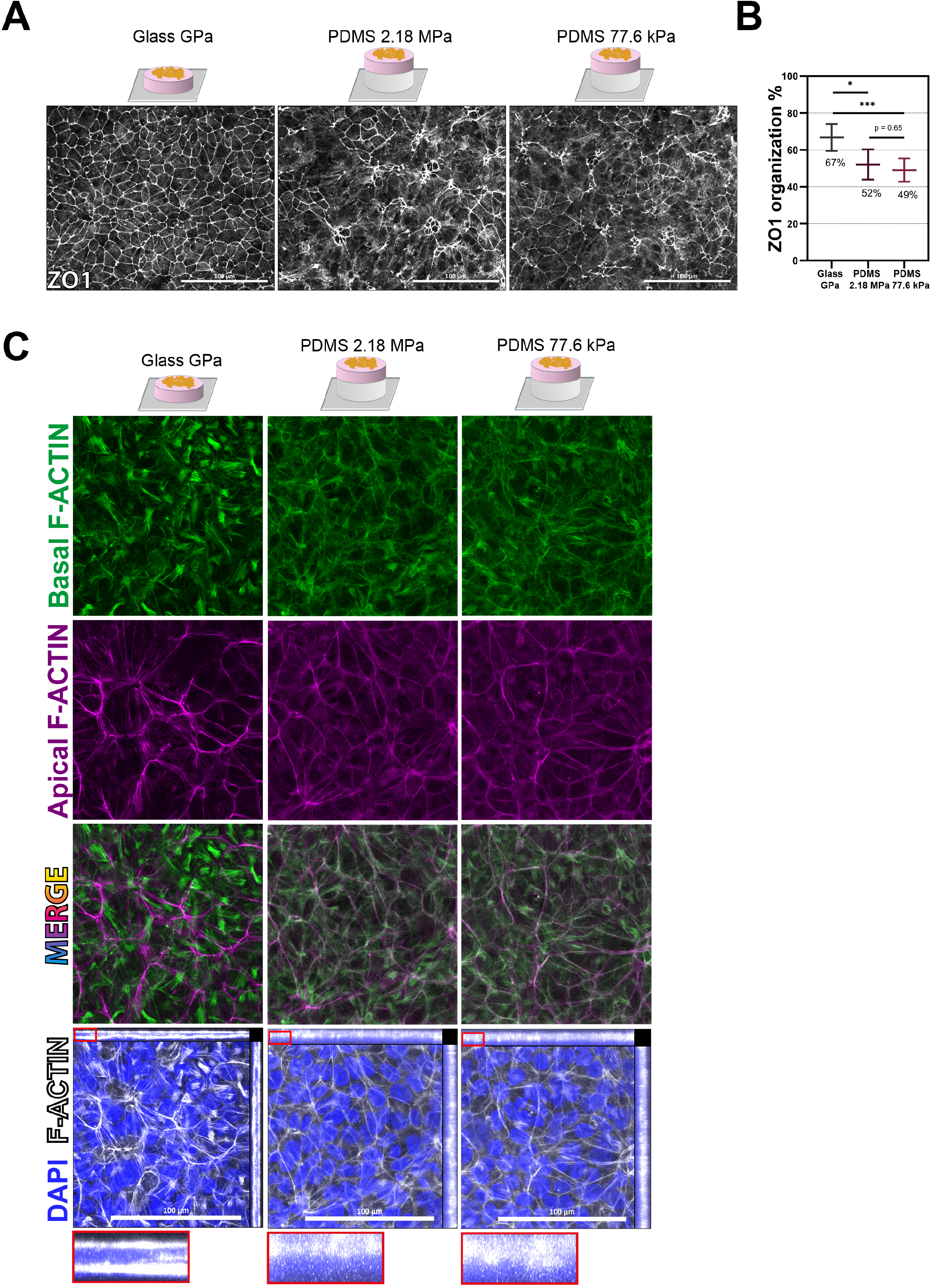
Epithelial and cytoskeletal organization of hiPSCs are perturbed on PDMS gel substrates. **(A)** Representative IF images of the tight junction protein ZO1 in hiPSCs grown on glass (left), 2.18 MPa PDMS gel (middle), and 77.6 kPa PDMS gel (right). Cells were fixed and images were taken 24 hours after the removal of ROCK inhibitor (48 hours after cell seeding at 110,000 cells/cm^2^). **(B)** ZO1 organization, defined as the percentage of the frame of view covered by ZO1-bounded cells, in the three stiffness conditions at the time point illustrated in **(A)**. Areas of ZO1-bounded cells were detected automatically. At least 3 randomly selected frames of view were taken per condition, covering >5.28 mm² in total. N = 6 ratio paired replicates. **(C)** Representative IF images of the nuclear marker DAPI and of phalloidin-labelled F-ACTIN in hiPSCs grown on glass (left), 2.18 MPa PDMS gel (middle), and 77.6 kPa PDMS gel (right). Cells were fixed and images were taken 24 hours after the removal of ROCK inhibitor (48 hours after cell seeding at 110,000 cells/cm^2^). Shown in green (first row) and magenta (second row) are the x-y planes of basally and apically located actin cortex respectively, detected automatically. In the bottom row (x-y, x-z, and y-z planes) and red insert (magnified x-z plane), maximum intensity projections of actin across all planes are shown. Scale bar in all images = 100 µm. Data are represented as mean ± SEM, and p values show the results of a two-tailed t test (*, < 0.05; **, < 0.01; ***, < 0.001).

Having demonstrated that epithelial integrity is reduced on both PDMS gels versus glass, we next sought to determine whether other chemical and physical aspects of the PDMS gels could cause this lag in hiPSC epithelial organization. As the PDMS gels used are naturally hydrophobic, we first tested the effect of differences in surface chemistry on the dynamics of cell organization. To this aim hiPSCs were plated on plasma-activated gels, in order to reverse the hydrophobicity of the PDMS surfaces and make them hydrophilic like glass, and immunostained for ZO1. Image analysis at 24 hours post-seeding revealed impaired TJ formation in hiPSCs plated on plasma-activated gels to a similar extent as non-activated gels (Figure S2B). Next, we investigated the potential impact of the gel surface roughness. Indeed, the PDMS gels’ thin layers are prepared via spin-coating of liquid droplets of base:catalyst mixtures, followed by an overnight incubation at 60°C to induce reticulation. This method is expected to produce a smooth gel surface topography similar to that of glass. To check whether some roughness may subsist that may impact cells’ ability to form contacts with one another, we estimated the PDMS gels’ surface topography by overlaying them with a dense layer of microscopic fluorescent beads (Figure S2C). A somewhat wider distribution in height of the beads was seen on the PDMS gels versus glass, especially with the 2.18 MPa PDMS gel. However, as the loss of epithelial integrity is similar for both PDMS gels (Figures 2B and S2C), the seemingly larger height of the bead layer on the PDMS gels (Figure S2C) is likely an artifact due to their different refractive indexes, We thus conclude that the PDMS gel surface topography is comparable to glass, with no impact on TJ formation. Finally, we tested whether there is any relationship between the substrate chemical composition and the epithelial integrity. Comparing glass with PDMS gel and PA hydrogel of stiffnesses of the same order (77.6 and 32 kPa, respectively), we found that a loss of epithelial integrity characterizes all gel types, demonstrating the phenotype of epithelial disorganization is independent of the gel chemistry (Figure S2D; percentage of surface covered by ZO1-bounded cells: glass 91%±1%; PA 32 kPa 82%±1%; PDMS 77.6 kPa 83%±3%). Taken together, these results point to substrate stiffness as the dominant factor in determining the TJ organization of the hiPSC epithelial layer without any significant contribution from gel surface chemistry, surface topography, or chemical composition.

As changes in cell-matrix and cell-cell contacts are known to remodel the cellular cytoskeleton^12^, we examined the distribution of the ACTIN cytoskeleton, namely the actin cortex which is made of actin filaments (F-ACTIN), in hiPSCs grown on the two PDMS gels of different stiffnesses versus hiPSCs grown on glass. Analysis of F-ACTIN localization demonstrated the occurrence of an ACTIN cytoskeleton remodeling in cells on PDMS gels when compared to glass (Figure 2C). In particular, on glass, there was a markedly increased presence of basal stress fibres when compared to PDMS gels, where concentrated actin fibers were mostly apico-lateral (Figure 2C). Z- projections of F-ACTIN on the three substrates further emphasized this remodeling (Figure 2C, red insert).

### Stiffness-mediated control of cell differentiation relies on epithelial organization

We next assessed whether disruption of the epithelial integrity in hiPSCs grown on gels could underlie their increased differentiation potential into mesendoderm and definitive endoderm. This was addressed by restoring the epithelial integrity in hiPSCs using lyso-phosphatidic acid (LPA), a small molecule known to accelerate epithelial organization and the formation of cell contacts through swelling of the apical membrane^35^. As shown in Figure 3A, a 24 hours treatment of hiPSCs grown on both PDMS gels with 5 µM LPA was sufficient to promote TJ organization and epithelial integrity of cells to levels comparable to those of cells grown on glass (almost fully organized following LPA treatment; Figure 3A). Consistent with an increase of TJ organization of hiPSCs on PDMS gels, their differentiation efficiency was reduced to the same level as that of hiPSCs on glass upon LPA treatment (Figure 3B), as estimated by the quantitative analysis of BRA-positive cells (Figure 3C; percentage of BRA-positive cells upon Activin treatment: glass 28.1%±5.8%; PDMS 2.18 MPa: 60.3%±10.4%; PDMS 77.6 kPa: 54.5%±11.9%; percentage of BRA-positive cells upon Activin and LPA treatment: glass: 2.4%±0.5%; PDMS 2.18: MPa 5.5%±2.6%; PDMS 77.6 kPa: 3.8%±1.8%). In conclusion, LPA treatment resulted into a drastic reduction of the differentiation efficiency of hiPSCs independently of substrate stiffness (nearly 0% in all conditions) thus highlighting a key role of epithelial organization in stiffness-mediated control of cell differentiation.

**Figure 3.**
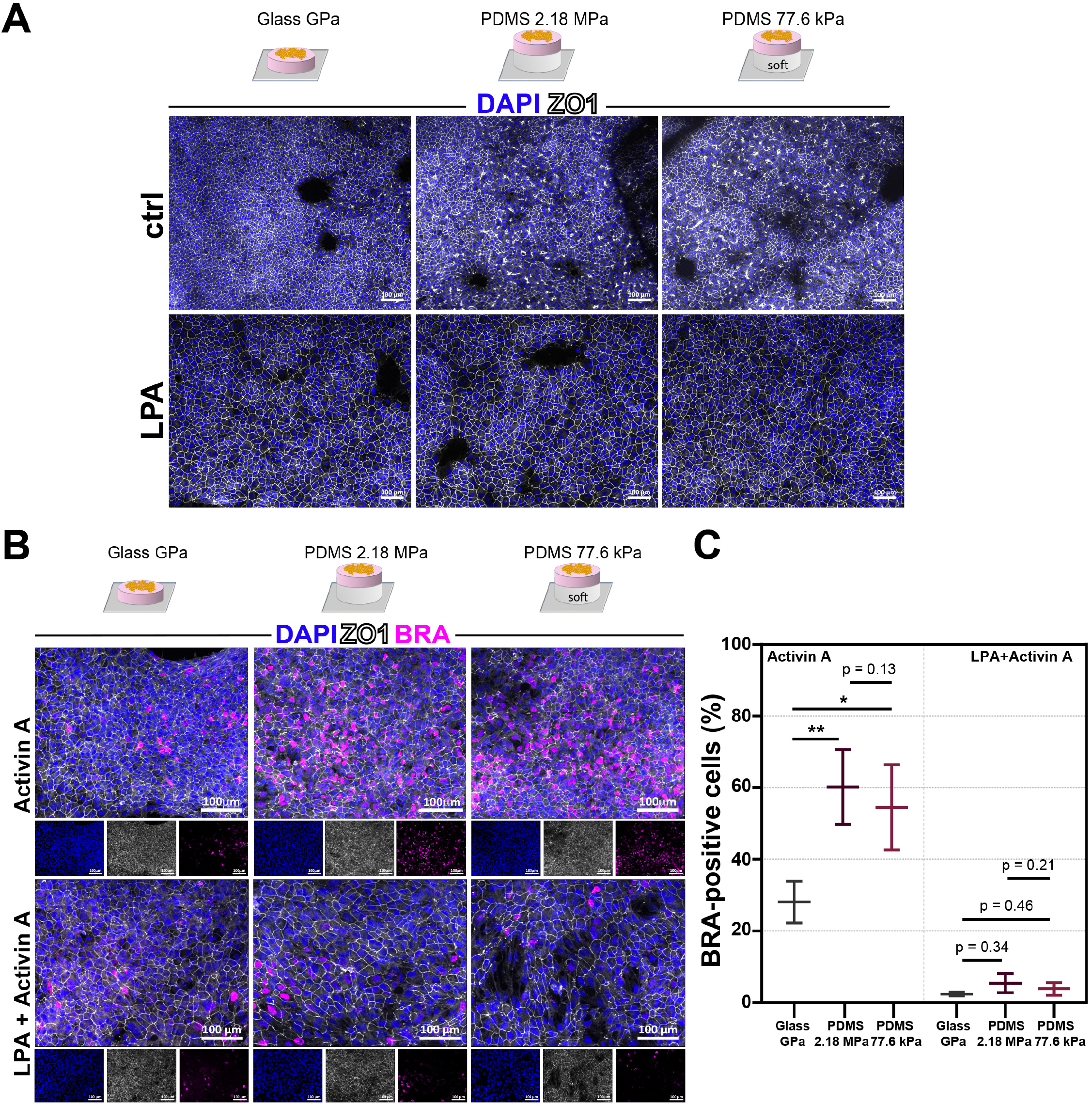
Substrate stiffness controls early lineage commitment in hiPSCs via a mechanism dependent on epithelial disorganization. **(A)** Representative IF images of the nuclear marker DAPI and of the tight junction protein ZO1 in hiPSCs grown on glass (left), 2.18 MPa stiffness PDMS (middle), and 77.6 kPa stiffness PDMS (right). 6 hours after cell seeding, ROCK inhibitor was removed and either blank (top) or 5 µM lyso- phosphatidic acid (LPA, bottom), was added to the culture media. Cells were fixed and images were taken 24 hours after the removal of ROCK inhibitor (30 hours after cell seeding at 130,000/cm^2^). **(B)** Representative IF images of DAPI, ZO1 and the of the mesendoderm marker BRA in hiPSCs grown on glass (left), 2.18 MPa stiffness PDMS (middle), and 77.6 kPa stiffness PDMS (right). Cells were fixed and images were taken 24 hours after the addition of 100 ng/ml Activin A (48 hours after the removal of ROCK inhibitor, addition or not of 5 µM LPA, and 72 hours after cell seeding at 50,000 cells/cm^2^). **(C)** Quantification of the number of BRA-positive cells in the three stiffness conditions at the time point illustrated in (B), in the absence (panels Activin A) and in the presence of LPA (panels LPA + Activin A). BRA-positive cells were defined as those whose mean fluorescence intensity was above the 95^th^ percentile of the mean BRA intensity in Activin A-negative control cells at the same time point and stiffness. At least 10 randomly selected frames of view were taken per condition, covering >2.23 mm² in total. N = 3 paired replicates. Scale bar in all images = 100 µm. Data are represented as mean ± SEM, and p values show the results of a two-tailed t test (*, < 0.05; **, < 0.01; ***, < 0.001).

### Ultra-soft silicone provides a reliable alternative to study hiPSC differentiation

Given that hiPSCs preferentially form 3D aggregates on ultra-soft PA hydrogels (< 10 kPa), we tested a different type of silicone gel, named QGel, more commonly used in electronics and much less-documented within biological sciences, whose stiffness can be tuned down to very low values, in the kPa range^36^. Utilizing the same method as for the former PDMS-based substrates, i.e. spin-coating base:catalyst mixture droplets of varying ratios onto glass coverslips, we fabricated QGel-based thin substrates with three stiffnesses of 31.2, 1.9, and below 1.9 kPa, which was the limit of detection for stiffness measurement (Table 1). We next tested whether hiPSCs were able to attach and grow on these ultra-soft QGels. In contrast to ultra-soft PA hydrogels with a Young’s modulus of 2.15 kPa, hiPSCs reliably formed a 2D epithelium on the QGel softer than 1.9 kPa (Figure 4A). Notably, cells cultured on ultra-soft QGels exhibited a similar perturbation of epithelial organization as seen on the softest PDMS gel (77.6 kPa) when compared to glass (Figures 4B and 4C; percentage of surface covered by ZO1-bounded cells: glass 49±7; PDMS 77.6 kPa 29%±4%; QGel <1.9 kPa 27%±5%).

Next, we tested the differentiation ability of hiPSCs on QGels by triggering endoderm lineage entry. As shown by the analysis of SOX17-positive cells, hiPSCs grown on QGels show increased differentiation efficiency when compared to hiPSCs grown on glass (Figure 4D and 4E). Moreover, differentiation level of hiPSCs on the softest QGel was comparable to that on the 77.6-kPa PDMS gel (Figure 4D and 4E; percentage of SOX17-positive cells: glass 4.2%±1.1%; PDMS 77.6 kPa 20.2%±2.7%; QGel <1.9 kPa 23.2%±0.4%) thus showing that QGels replicate loss of TJ organization and increased differentiation efficiency of hiPSCs on substrates softer than glass. In conclusion, our results show that QGel ultra-soft matrices are compatible with expansion and differentiation of hiPSCs. In this way QGels provide a new substrate to test the effects of ultra-soft gels with stiffnesses lower than 10 kPa for cell lines, like some hiPSC lines, which are not compatible with ultra-soft PA hydrogel.

**Figure 4.**
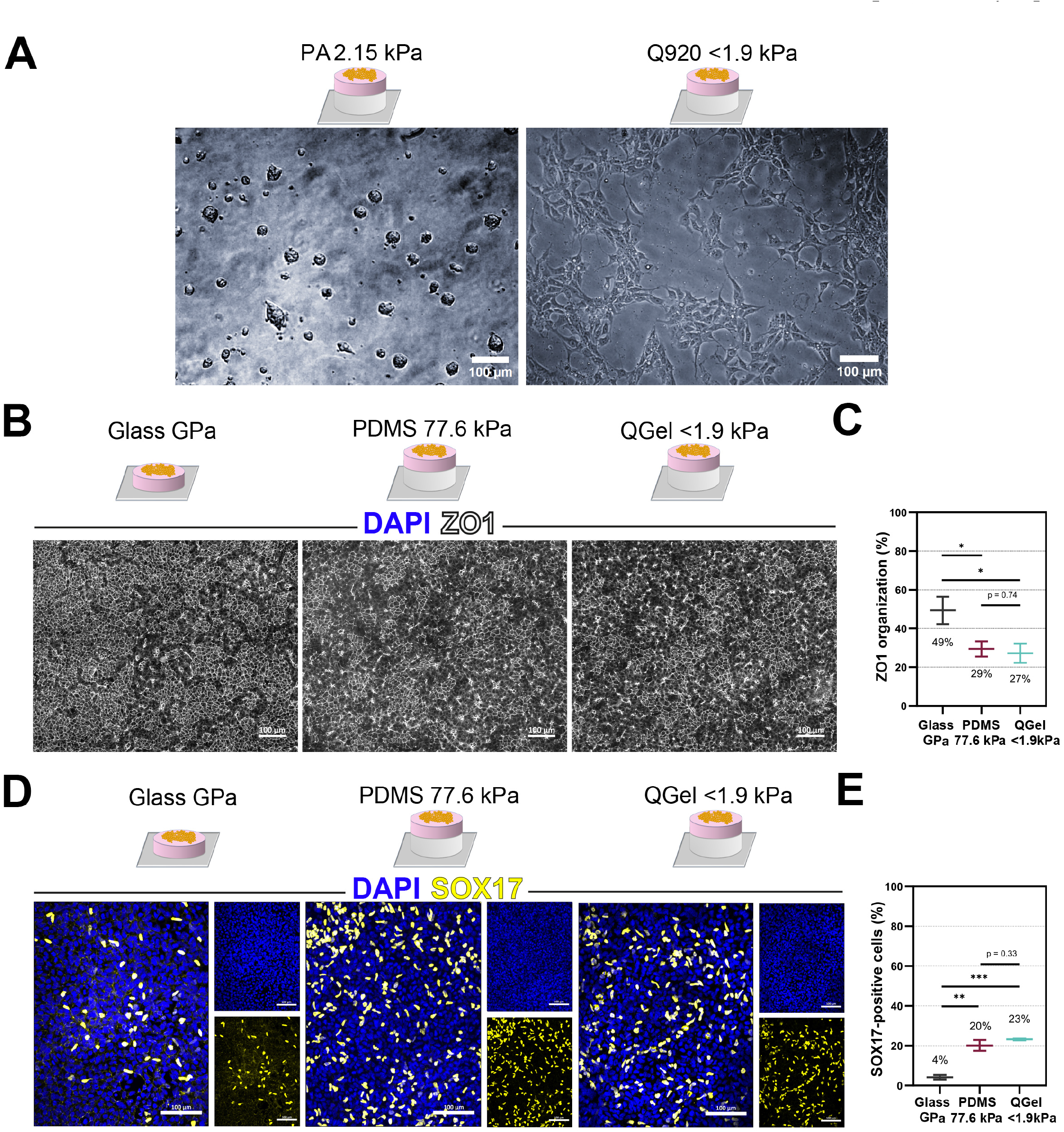
QGels permit expansion and differentiation of hiPSCs on ultra-soft substrates. **(A)** Bright field images of the inability/ability of hiPSCs to adhere and grow as a 2D epithelium on PA hydrogels (2.15 kPa, left) and on QGel (<1.9 kPa, right). Cells were fixed and images were taken 24 h after cell seeding at 110,000 cells/cm^2^. **(B)** Representative IF images of the tight junction protein ZO1 in hiPSCs grown on glass (left), soft PDMS gel (77.6 kPa, middle), and ultra-soft QGel (<1.9 kPa, right). Cells were fixed and images were taken 24 hours after the removal of ROCK inhibitor (48 hours after cell seeding at 110,000 cells/cm^2^). **(C)** The ZO1 organization, defined as the percentage of the frame of view covered by ZO1- bounded cells, in the three stiffness conditions at the time point illustrated in **(B)**. Areas of ZO1- bounded cells were detected automatically. At least 4 randomly selected frames of view were taken per condition, covering >7.04 mm² in total. N = 4 unpaired replicates. Note that hiPSCs plated on glass displayed approximately 50% to 70% of TJ organization in different experiments depending on the growth rate of cells at the plating stage (compare Figures 4C and 2B). **(D)** Representative IF images of the nuclear marker DAPI and of the definitive endoderm marker SOX17 in hiPSCs grown on glass (left), 77.6 kPa stiffness PDMS (middle), and <1.9 kPa stiffness QGel (right). Cells were fixed and images were taken 24 hours after the addition of 0.2% FBS (48 hours after the addition of 100 ng/ml Activin A, 72 hours after the removal of ROCK inhibitor, and 96 hours after cell seeding at 50,000 cells/cm^2^). **(E)** Quantification of the number of SOX17-positive cells in each of the three stiffness conditions at the time point illustrated in (D). SOX17-positive cells were defined via automatic thresholding. At least 3 randomly selected frames of view were taken per condition, covering >21.2 mm² in total. N = 3 unpaired replicates. Scale bar in all images = 100 µm. Data are represented as mean ± SEM, and p values show the results of a two-tailed t test (*, < 0.05; **, < 0.01; ***, < 0.001).

### Epithelial structures point to an increased fluidity of hiPSC colonies on softer substrates

Having found that cells grown on PDMS/QGel silicone gels and PA hydrogels show a disruption of epithelial integrity versus cells grown on glass, we next sought to decipher whether this phenotype correlates with changes of hiPSC physical properties in response to matrix stiffness. We performed live imaging of cells growing on the 77.6 kPa PDMS gel and glass and examined both single cells and colonies, 0-6 h and 24-30 h after and the removal of ROCK inhibitor (Videos S1 to S4). We measured the correlation coefficient between frames from these time lapses. This coefficient quantifies how different a video frame is compared to the preceding one, i.e. how much cells have moved between the two frames. This approach allowed us to quantify how the motility of hiPSCs changes on gel substrates compared to glass, both outside (single cells) and inside of colonies^37–39^. At both time windows, cells growing on the PDMS gel, either as single cells or colonies, are more motile when compared to cells growing on glass. Indeed, though very close, the correlation coefficients are significantly different between glass and PDMS (correlation coefficient 0-6 h: glass GPa 0.8556±0.0013, PDMS 77.6 kPa 0.8471±0.0022; correlation coefficient 24-30 h: glass GPa 0.8944±0.0011, PDMS 77.6 kPa 0.8862±0.0009; Figure 5A). The finding that both single cells and cells within colonies are more motile on the PDMS gel may explain the impaired epithelial integrity we observed compared to glass: more motile cells on soft substrates may require more time to form stable TJs and reach a high level of epithelial organization.

**Figure 5.**
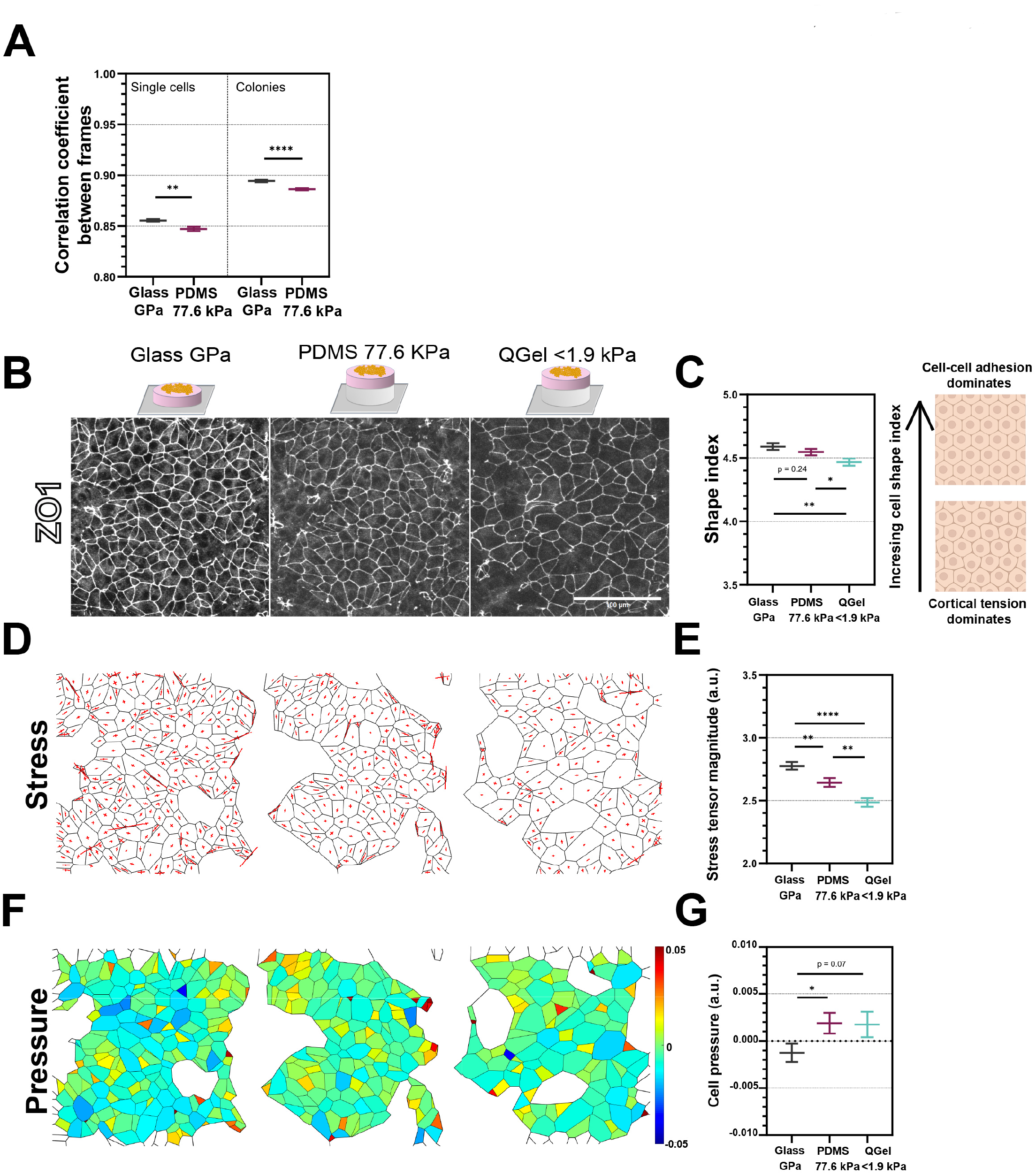
HiPSCs growing on gel substrates are more motile and display lower stress and higher cytoplasmic pressure. **(A)** Quantification showing the correlation coefficients between successive frames of the videos shown in Supplemental Movies 1-4. An automatic mask was used to remove non-cell areas from each frame and arrays representing the frames were subjected to a 2D correlation analysis. Frames were taken with a 5 min time interval over 6 h, giving N = 72 pairs of successive frames in each condition. Data are represented as mean ± SEM, and p values show the results of a two- tailed, unpaired t test (*, < 0.05; **, < 0.01; ***, < 0.001). **(B)** Magnified regions of epithelia taken from the data quantified in Figure 4C, showing hiPSCs grown on glass (left), soft PDMS gel (77.6 kPa, middle), and ultra-soft QGel (<1.9 kPa, right) with similar percentages of ZO1 organization. Scale bar = 100 µm. **(C)** (Left) quantification of the shape index, defined as the cell perimeter divided by the square root of the cell area, in each of the three images in **(B)**. Cells are segmented automatically using Tissue Analyzer^63^. Data are represented as mean ± SEM, and p values show the results of a two- tailed unpaired t test comparing N≥135 cells (* < 0.05, ** < 0.01, *** < 0.001). (Right) illustration of the epithelial phenotype of decreasing shape index, which is inversely proportional to cell circularity. **(D)** Maps of the stress tensors across all cells in each of the three images in (B) predicted by Bayesian force inference^40^. Red crosses in the centers of segmented cells are sized relative to the magnitudes (arbitrary units) of stress tensors parallel and perpendicular to the axis of maximum stress. **(E)** Quantification of the stress tensors (arbitrary units) shown in the map in (D), taking the tensor value along the axis of maximum stress for each cell. Data are represented as mean ± SEM, and p values show the results of a two-tailed unpaired t test comparing N≥135 cells (* < 0.05, ** < 0.01, *** < 0.001). **(F)** Maps of the pressures (normalized to an average pressure of 0)^43^ of all cells in each of the three images in (B) predicted by Bayesian force inference. Analogous to cell stress in the z direction, the predicted pressure of each cell maintains its current size. **(G)** Quantification of the cell pressures (arbitrary units, normalized to an average pressure of 0)^43^ shown in the map in (F), taking the tensor value along the axis of maximum stress for each cell. Data are represented as mean ± SEM, and p values show the results of a two-tailed unpaired t test comparing N≥135 cells (* < 0.05, ** < 0.01, *** < 0.001).

Next, we analyzed how the epithelial structure of hiPSCs differs on substrates of different stiffnesses (glass vs soft PDMS 776.6 kPa vs ultra-soft QGel <1.9 kPa; Figure 4B). To do so, we applied a Bayesian force inference method^40^, which is well-used and experimentally validated in studies of epithelial force dynamics^41–43^. Based on the assumption that forces by cell-cell interfacial tensions and intra-cellular pressures balance at each vertex (i.e. the junction point between at least 3 cells) this method predicts various epithelium features including the magnitude of global interfacial stress experienced by cells across the epithelium, and the intra-cellular pressure each cell needs to produce to maintain its shape. Taking regions of epithelia with representative areas of organization in the three stiffness conditions at 24 hours following ROCK inhibitor removal (Figure 5B), we first analyzed the shape index of cells in the three epithelia. The shape index, defined as the cell perimeter divided by the square root of the cell area, illustrates the competition between cell cortical tension and cell-cell contacts that both affect the cell shape. In cell layers with strong epithelial organization, a lower shape index indicates that cortical tension dominates the cell shape, whereas a higher shape index indicates that cell-cell adhesion has a stronger influence on cell shape^38^. In our study, hiPSC layers are more disorganized and we used the shape index to estimate differences in cell-cell adhesion at the time point used to trigger differentiation. We found that the shape index of cells progressively decreased with substrate stiffness (glass: 4.59±0.025; PDMS 4.546±0.026; QGel 4.467±0.028; Figure 5C), suggesting stronger and more influential cell-cell contacts on stiffer substrates for our hiPSC structures. Next, the Bayesian force inference was applied to the same images to estimate the interfacial stress and the intra-cellular pressure. As cells from the same batch were seeded at the same density on the three substrates and observed at the same time point, we made the assumption that the three samples are initially similar and that the differences in stress and pressure arise uniquely from behavioural differences due to the substrates. This approach provided evidence that cells on softer substrates, compared to glass, likely experience lower stress (Figure 5D and 5E; glass: 2.78±0.03, PDMS: 2.64±0.04, QGel: 2.49±0.03) and higher cytoplasmic pressure (Figure 5F and 5G; glass: −0.0013±0.001, PDMS: 0.0019±0.0011 QGel: 0.0017±0.0014), parameters that increase the cells’ ability to change their shape and size.

Together, the results of the live imaging analysis and Bayesian force inference suggest that the epithelial organization of cells within colonies on stiffer substrates is less fluid, due to lower cell motility, and that cells are tightly connected with higher interfacial tension. In contrast, on softer substrates, the increased motility of cells leads to more fluid colonies with less cell-cell contacts and, consequently, a slower rate of epithelial organization.

## Discussion

Understanding the mechanisms behind mammalian lineage commitment remains a considerable challenge, especially in human embryos. The prevailing idea is that crosstalk between biochemical and physical cues regulates the early processes of embryonic development. HPSCs, serving as *in vitro* models of the human epiblast, offer a unique opportunity to dissect these intricate processes while permitting independent manipulation of the underlying mechanisms. In the present study, we have explored whether the competence of hiPSCs to undergo mesendoderm and definitive endoderm differentiation in response to Activin A stimulation can be modulated by altering substrate stiffness as the selected physical cue. We demonstrated that differentiation of hiPSCs along these lineages can be improved when hiPSCs are cultured on substrates softer than glass, such as PDMS/QGel silicone gels and PA hydrogels with stiffnesses ranging between a few MPa and a few kPa, which are far below the GPa stiffness of glass. Of note, hiPSCs cultured on these gels exhibit alterations in TJ formation and a disrupted epithelial integrity. Furthermore, cells are more motile and exhibit alterations in shape, experience less forces and generate higher internal pressures. These phenotypes cause a delay in TJ formation kinetics, which is the basis of the increased differentiation potential of hiPSCs on gels, as demonstrated by rescue experiments. Thus, the results of this study provide a mechanistic insight into how physical inputs from the microenvironment modify the epithelial properties of hiPSCs, which ultimately determine their response to differentiation triggering biochemical signals.

Our temporal observation of TJ formation shows that substrates softer than glass do not alter the molecular mechanisms involved in TJ assembly, but instead regulate the kinetics of TJ formation by causing a delay in the stabilization of these cell-cell contacts after cell dissociation and seeding. Our *in silico* analysis indicates that the hiPSC layer acquires distinct epithelial mechanical properties on gels, displaying weaker and less influential cell-cell contacts, as well as increased fluidity within colonies. Furthermore, our live imaging analysis demonstrates that cells within hiPSC colonies exhibit higher motility levels, consistent with the increased cell layer fluidity. Therefore, it is probable that the delay in TJ formation on gels results from the instability of cell position within colonies, leading to an extended period required to establish stable cell-cell contacts. Future work on signal perception in epithelial tissues should seek to further quantify how the mechanoenvironment affects the fluidity of the cell layer.

Recent research indicates that the TJ scaffolding protein ZO1, which connects the TJ transmembrane proteins to the cytoskeleton^44^, plays a role in regulating TJ assembly that is in part dependent on tension^45^. Notably, the mechanical forces acting on ZO1 depend on extracellular matrix stiffness^46^. This implies that ZO1 may help create junctions by adapting the cytoskeletal forces generated in the cell body to the developing TJs. In support of this are our results showing broad cytoskeletal remodeling on substrates softer than glass, and cell shape analysis showing tension generated by the actin cortex becoming more dominant over forces from TJs. Based on our findings, substrate stiffness has an impact on the kinetics of TJ formation. Further research is required to determine whether these changes are associated with specific mechanical tension on ZO1.

Through our rescue experiments, we present evidence that the kinetics of TJ formation plays a crucial role in the regulation of hiPSC fate acquisition in response to differentiation factors. As per our previous findings, the loss of morphogen regulator GLYPICAN-4 in hiPSCs causes disruption in epithelial integrity with areas of TJ formation being affected^35^. This phenotype results in the Activin A receptors being exposed to the culture medium, thereby enhancing hiPSCs’ capacity to detect Activin A and maintain activation of the Activin A pathway over a prolonged period. This situation results in an enhanced differentiation potential into mesendoderm and definitive endoderm lineages. Based on these results, we hypothesize that hiPSCs on gels develop greater competence in perceiving the Activin A differentiation signal due to areas of disrupted TJs in which receptors might be more exposed, resulting in more efficient differentiation. Additionally, it is also likely that a modification in fluidity of the hiPSC colonies on gels (as mentioned earlier) increases Activin A receptor accessibility across the entire hiPSC layer, leading to improved Activin A signal perception. Crucially, our rescue of epithelial organization ceased differentiation in response to Activin A almost entirely in all conditions. This demonstrates that the stiffness-mediated effect on cells’ sensitivity to Activin A signaling depends on the effects of stiffness on epithelial organization.

Interestingly, this disruption of epithelial integrity for hiPSCs grown on gels does not perturb self-renewal or the expression of pluripotency genes, nor promotes premature expression of mesendoderm, endoderm or other lineage markers (present study and in^35, 47^). Thus, substrate stiffness-dependent alterations in TJs alone do not appear sufficient to promote differentiation or determine lineage fate choices. The observed maintenance of stemness/pluripotency of hiPSCs grown on gels is consistent with previous studies from Przybyla et al. and Guo et al. showing that changes in the PA hydrogel stiffness ranging from 60 kPa to 0.4 kPa, and 3.5 kPa, respectively, do not affect these biological properties of hESCs^12, 15^. Notably, Guo et al. found that hESCs grown on a 0.4-kPa PA hydrogel underwent a significant change in shape, resulting in colonies with a rounded morphology similar to those we observed in our study with hiPSCs grown on PA hydrogels with stiffnesses lower than 10 kPa^15^. Contrary to the previously mentioned outcomes, Musah et al. discovered that utilizing a soft PA hydrogel with a Young’s modulus around 0.7 kPa causes down regulation of pluripotency genes while promoting neuronal lineage entry and differentiation into post mitotic neurons, even if the cells are grown in a self-renewing medium^48^. Similarly, Chen et al. demonstrated that using soft PA hydrogels with stiffnesses of 3, 15, and 33 kPa causes down regulation of pluripotency genes and primes hiPSC differentiation towards the endoderm lineage, without requiring endoderm induction medium^13^. Unfortunately, it was not reported in these two studies if the hPSCs grown on these soft PA-based substrates maintain their epithelial morphology or develop a spherical colony shape, which could impact their behavior. While designing substrates to bypass the need for differentiation-inducing factors is fascinating, the aforementioned studies also suggest that different hPSC lines could react differentially to substrate stiffness. In particular, on soft PA-based substrates with similar stiffnesses (0.4 kPa in^12^, and around 0.7 kPa in^48^), hPSCs can retain stemness and pluripotency^12^ or develop into neuronal lineages^48^. These findings also suggest that additional unknown factors present in culture conditions might cooperate with substrate stiffness in the establishment of hPSC behavior, which could have implications for a reproducible lineage acquisition and maintenance. We suggest that identifying the optimal substrate stiffness to preserve pluripotency in hPSCs, while enhancing differentiation potential under differentiation conditions, could prevent such variations. This would enable precise control of culture conditions for cell therapy, disease modeling, and bio-mechanistic studies.

Our research here and others’ findings indicate that PA hydrogels with stiffness lower than 10 kPa may not always support the maintenance of hiPSC epithelial morphology. A common reasoning is that cells will preferentially form 3D aggregates and detach (present studies and^33, 34^). Our experiments, using QGels of varying stiffness down to less than 2 kPa, indicate that this kind of substrate should enable the maintenance of self-renewal and pluripotency of hPSCs for which ultra-soft PA hydrogel are not compatible for growth and maintenance in an undifferentiated state. The QGel silicone gels are frequently utilized in electronics as they provide protection against moisture, vibration, thermal or mechanical shock. While the material has been used sporadically as an ultra-soft substrate to study cell mechanobiology^36, 49–51^, here we present for the first time its suitability for hPSC culture and studies of cell differentiation in physiological stiffness conditions. Our findings reveal that silicone QGels are innovative materials that allow for the culture and differentiation of hPSCs, without inducing any adverse effects, when combined with bioactive molecules like Matrigel.

Additionally, QGels seem to offer some technical advantages compared to PA hydrogel.

Traditional PA hydrogel layers are most commonly fabricated by sandwiching the liquid gel mixture between glass surfaces^52^ however silicone gels are sufficiently viscous to enable droplet spin-coating onto glass coverslips, which allows for more precise and reliable control of gel properties such as thickness and surface smoothness. Further, the dependence of PA hydrogels on initiators (ammonium persulfate and TEMED) introduces batch variance in gel polymer lengths and elasticity https://www.bio-rad.com/webroot/web/pdf/lsr/literature/Bulletin_1156.pdf. It has also been shown that cells are less able to form stable focal adhesions on ultra-soft PA hydrogels, due to these being more porous than silicone gels^53, 54^. We thus hypothesize that the adhesion, buffering, and elasticity features of QGel silicone gels facilitate the adhesion and growth of hPSCs, which are otherwise unsuitable for culture on ultra-soft but porous substrates like PA hydrogels.

In conclusion, our work highlights that, by optimizing the mechanical properties of the extracellular substrate, hPSCs can be maintained in a self-renewal pluripotent state, and mesendoderm as well as definitive endoderm differentiation can be enhanced. We also provide evidence of the mechanism underlying the enhanced differentiation potential of hiPSCs grown on gels by showing that substrate stiffness-triggered changes in the epithelial cell layer architecture can facilitate sensing of signaling proteins. We expect that this mechanism will be of more general relevance in other hPSC lineage entry processes and differentiation events. The interdisciplinary nature of our research provides novel perspectives on how cells combine physical and biological cues to initiate crucial processes, emphasizing the significance of investigating the function of composite interfaces in development, physiology, and pathology.

## Limitations of the study

In this study, our focus was on the impact of matrix stiffness on differentiation of hiPSCs into mesendoderm and endoderm lineages, and we revealed the underlying cellular mechanisms that drive this differentiation. It remains to be verified if TJ disruption, triggered by decrease of the matrix stiffness, also affects neuroectoderm and mesoderm differentiation, which will be explored in the future.

Bayesian force inference was used to analyze the mechanical differences between representative epithelial structures after 48 hours on different substrates. While it would be of interest to analyze these mechanical properties in different contexts, for example over larger areas or at different stages of epithelial organization, these experiments are currently subject to both hardware and software limitations. The analysis was also done assuming that hiPSC epithelial organization on different substrates could be directly compared. While outside the scope of this study, the use of a gradient stiffness gel would facilitate direct comparisons of epithelial forces across different stiffnesses.

## STAR Methods

### Resource availability Lead contact

Further information and requests for resources and reagents should be directed to and will be fulfilled by the lead contact, Dr. Rosanna Dono

### Materials availability

This study did not generate new unique reagents.

### Data and code availability

Data: All data reported in this paper will be shared by the lead contact upon request.

Code: This paper does not report original code.

Any additional information required to reanalyze the data reported in this paper is available from the lead contact upon request.

### Experimental model and study participant details

WT 029 hiPSCs were courtesy of Michael Kyba^55^. HiPSCs were maintained on tissue culture-treated plastic, coated with Matrigel (BV 354277, Corning) according to supplier recommendations, in mTeSR1 medium (85850, Stemcell Technologies) supplemented with 1% Penicillin/Streptomycin cocktail (15140122, Invitrogen), and maintained in a humidified 37°C incubator with 5% CO_2_. Medium was changed every 24 hours, and cell cultures were dissociated using Accumax (SCR006, Millipore) and passaged 1/10 ad-hoc prior to confluency, approximately every 3-4 days. Media were supplemented with 10 µM ROCK inhibitor (1254, Tocris) in the 24 hours following passage to facilitate cell adhesion and survival.

## Method details

### Gel fabrication

PDMS and QGel silicones (Sylgard 184, Dow Corning; QGel 920, CHT Silicones) were both fabricated by mixing each base with its respective cross-linker solution (TABLE 1). For each gel, the two solutions were mixed and degassed for 15 min. 100 µL of the mixture was spin-coated onto #1.5 round coverslips (170 µm in thickness, 13 mm in diameter) at 1,200 rpm for 60 s. Gels were cured, i.e. reticulated, by overnight incubation at 60 °C. Where appropriate, gel hydrophobicity was reversed to hydrophilicity via a 30-s plasma treatment.

PA hydrogels were prepared by adapting previously described protocols^56^. Gel binding on the same glass coverslips as for PDMS gels was facilitated by activating the glass surface using a 1:1:14 solution of acetic acid (695092, Sigma)/bind-silane (10700467, VWR)/ethanol. The coverslips were washed twice with ethanol and air-dried for 10 min. Acrylamide and bis-acrylamide (1010140/1610142, Bio-Rad) were mixed in phosphate-buffered saline (PBS) at ratios corresponding to the desired stiffnesses (TABLE 1). The polymerization reaction was catalyzed by adding 0.5% ammonium persulfate (A3678, Sigma) and 0.1% TEMED (T9281, Sigma). Before polymerization could occur, 10-µL drops were quickly added (coated with hydrophobic RAIN-X) and the drops were covered with the activated coverslips. PA hydrogel were left to polymerize for ∼30 min, then washed three times with PBS. Hydrogels were then activated to facilitate cell adherence by 10-min incubation with 0.5 mg/mL Sulfo-SANPAH (22589, ThermoFisher Scientific) in MilliQ water under 365-nm UV light. After activation, gels were washed three times with PBS.

Prior to cell seeding, all substrates (including glass) were functionalized using Matrigel (BV 354277, Corning) according to manufacturer recommendation.

The surface roughness of substrates was tested by overlaying a solution of fluorescent beads (1 µm in diameter, F8816, ThermoFisher Scientific). 3.2 µL of the bead stock solution was mixed in 500 µL Matrigel and overlaid onto glass or gel-coated coverslips for 20 min at room temperature, followed by three PBS washes.

The Young’s moduli and standard deviations are respectively the weighted average and weighted standard deviations calculated from the N measurements (the inverse variance weighting was used to minimize the uncertainty). Note that we were not able to measure the softest QGel, but we expect from its composition to be softer than the 1.9 kPa QGel.

### Atomic force microscopy

Nanoindentation experiments were performed using Atomic Force Microscopy (AFM, NTEGRA- NTMDT) in contact mode, by controlling the displacement of the piezoelectric element in the direction perpendicular to the surface and the time rate required to perform this displacement. The deflection of the cantilever was continuously measured during loading and unloading from the sample surface. In all experiments, the same type of probe was used: CONT-Silicon –SPM- Sensor with a colloidal particle of diameter 6.62 μm±10% as indicated by the manufacturer (Nanosensors). The probe’s cantilever stiffness was measured using the Sader method^57^. Cantilever parameters were measured experimentally using the AFM setup (Resonance frequency and Quality factor) and optical microscopy (Width and Length). A total of four probes were used in the whole set of experiments with their characteristic parameters indicated in Table 2.

**Table 2:** Cantilever parameters of AFM probes.

| Probe | Resonance frequency [kHz] | Quality factor | Width [$\mu\text{m}$] | Length [$\mu\text{m}$] | Stiffness [N/m] |
| --- | --- | --- | --- | --- | --- |
| 1 | 14.25 | 61.56 | 47.6 | 454 | 0.217 |
| 2 | 18.64 | 84.96 | 47.6 | 450 | 0.417 |
| 3 | 18.63 | 88.90 | 46.0 | 448 | 0.437 |
| 4 | 18.60 | 78.32 | 49.3 | 450 | 0.413 |

Prior to experiments, AFM probes were cleaned in acetone, and exposed for 30 s to oxygen plasma. To reduce a strong attractive interaction between the tip and the samples, the tips were functionalized with trimethoxymethylsilane, making their surface hydrophobic. Similarly, all experiments were performed in water to avoid capillary forces. For each set of measurements, cantilever sensitivity, i.e. deflection versus piezoelectric element z-displacement, was determined by performing a force-distance measurement on a hard silicon sample and further used to normalize the data curves. Indentation measurements were performed at a rate 1 Hz and a maximum loading force of 10 nN was applied during loading.

### Extraction of Young’s modulus

Young’s moduli were extracted using the Oliver & Pharr method^58, 59^, which is suitable for processing both purely elastic and viscoelastic behaviors. The Young’s modulus is extracted from the value of the slope (S) at the beginning of the unloading curve. For an indenter with a spherical shape, the relation between S, the maximum force (F), the indentation depth (h), the final depth (after the indenter is fully unloaded) (h_f_), and the indentation modulus (M) is expressed as:

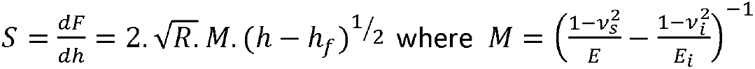

where E and E_i_ are the Young’s modulus of the sample and the indenter respectively, and ν_s_ and ν_i_, the sample and indenter Poisson’s coefficients respectively. E_i_ was fixed at 150 MPa, ν_i_ at 0.3 and ν_s_ at 0.5.

### Adherence, pluripotency, organization, and differentiation assays

For cell adherence and organization experiments, hiPSCs were seeded at a density of 110,000 cells/cm^2^ onto Matrigel-coated coverslips in mTeSR1 medium supplemented as described earlier. After 24 h, fresh media was added with the omission of ROCK inhibitor. After a further 24 hours (48 hours in total post-seeding), bright field images were taken for adherence assays, or cells were fixed for 10 min using 4% paraformaldehyde (PFA) in PBS and analyzed by immunocytochemistry for pluripotency and organization assays.

Differentiation experiments of hiPSCs into mesendoderm and definitive endoderm lineages were performed by following published protocols^25^. HiPSCs were seeded onto Matrigel-coated coverslips in mTeSR1 medium supplemented as described earlier, at densities of 110,000 and 50,000 cells/cm^2^ for mesendoderm and definitive endoderm differentiation experiments, respectively. After 24 hours, fresh medium was added with the omission of ROCK inhibitor. After a further 24 hours (48 hours in total post-seeding), medium was removed and cells were washed twice with RPMI medium (Invitrogen, ref. 21875034), followed by the addition of RPMI medium supplemented with 100 ng/mL Activin A (338-AC, R&D), while control groups were maintained in supplemented mTeSR1 medium. After a further 24 h (72 h in total post-seeding), cells were either fixed to analyze mesendoderm differentiation by immunocytochemistry, or further cultured in fresh RPMI supplemented with 100 ng/mL Activin A and 0.2% defined, heat-inactivated fetal bovine serum (FBS, SH30071.02E, Hyclone). After a further 24 hours (96 hours in total post-seeding), cells were fixed to analyze definitive endoderm differentiation by immunocytochemistry.

### Chemical rescue of epithelial integrity

For epithelial integrity rescue experiments, hiPSCs were seeded at 130,000 cells/cm^2^ density on Matrigel-coated coverslips in mTeSR1 medium supplemented with ROCK inhibitor. After 6 h, ROCK inhibitor was removed and either control medium or medium supplemented with 5 µM of LPA was added. After a further 24 hours (30 hours in total post-seeding), cells were fixed and analyzed by immunocytochemistry.

For differentiation experiments in rescued epithelia, hiPSCs were seeded at 110,000 cells/cm^2^ density on Matrigel-coated coverslips in mTeSR1 medium supplemented with ROCK inhibitor. After 24 hours, ROCK inhibitor was removed and media supplemented with 5 µM of LPA was added. After a further 24 hours (48 hours in total post-seeding), cells were fixed to analyze mesendoderm differentiation by immunocytochemistry.

### Immunocytochemistry

Following fixation, cells were washed three times with PBS. Cells were permeabilized in 0.3% Triton-x100 (108603, Millipore) in PBS for 10 min at room temperature and blocked with 3% bovine serum albumin (A9647, Sigma), 2% donkey serum (ab7475, Abcam), 0.3% Triton-x100 in PBS for 1 hours at room temperature. Cells were incubated with primary antibodies (Table 3) in blocking solution at 4°C overnight, followed by four PBS washes. Cells were then incubated with appropriate secondary antibodies (1/500 Invitrogen) in blocking solution with DAPI (D3571, Invitrogen) and Phalloidin-iFluor647 (where applicable, ab176759, ABCAM) for 1 hour at room temperature. Cells were then washed four times using 0.3% Triton-x100 in PBS, following which coverslips were rinsed twice in MilliQ water before mounting onto 1-mm thick glass slides using anti-fade mounting medium (P36930, Invitrogen).

**Table 3.** List of Primaries antibodies used for Immunocytochemical analyses.

| **Antigen** | **Specie** | **Company** | **Reference** | **Dilution** |
| --- | --- | --- | --- | --- |
| BRACHYURY | Goat | R&D | 967332 | 1/80 |
| E-CADHERIN | Rabbit | Cell Signalling | 3195 | 1/200 |
| NANOG | Rabbit | Cell Signalling | 4903 | 1/200 |
| OCT4 | Rabbit | SantaCruz | 5272 | 1/400 |
| SOX2 | Mouse | SantaCruz | sc365823 | 1/1000 |
| SOX17 | Goat | R&D | 967330 | 1/80 |
| ZO-1 | Mouse | Invitrogen | 33-9100 | 1/500 |

### Microscopy

Bright field images were taken on a Nikon Eclipse TS100 microscope using a DinoEye Edge AM7025X camera. Immunofluorescence images were acquired with either a Zeiss Axio Imager M2 wide-field or a Zeiss LSM 880 confocal microscope, using ZEN (version 3.4). Live cell imaging was carried out using a Zeiss Axio Observer Z1 fitted with a custom-built 37 °C 5% CO_2_ chamber. Cells were observed using a 20x objective over 6-h time periods and with a 5-min interval between frames to balance maximizing data acquisition and resolution with minimizing phototoxic exposure in order to ensure authenticity of cell behavior.

## Quantification and statistical analysis

### Immunofluorescence quantification

For quantification of BRA-positive mesendoderm cells, the number of positive cells showing nuclear localization of the transcription factor BRA were automatically counted by the FIJI software^60^ (version 2.14.0). BRA-positive cells were defined as those whose mean fluorescence intensity was above the 95^th^ percentile of the mean BRA intensity in Activin A-negative control cells at the same time point and stiffness. DAPI staining was used to define nuclear areas and total cell numbers using the StarDist^61^of the FIJI software. SOX17-positive definitive endoderm cells were identified using the FIJI plugin StarDist as above.

### Organization quantification

The Cellpose algorithm^62^ (version 2.0) was used to identify cells fully bound by ZO1 contacts in ZO1 immunofluorescent images. Connected regions of fully ZO1-bound cells were classified as areas of epithelial organization. The percentage organization in each image was calculated as the percentage of the total frame of view covered by areas of epithelial organization^35^ .

### Growth analysis

The confluency of cells in phase-contrast frames of time lapses was calculated using FIJI. Images were converted to binary images using automatic thresholds, and objects filtered according to a size minimum of 5000 px² (image calibration: 0.631 µm/px). The confluency at each time point was defined as the percentage of the frame of view taken up by objects.

### Motility analysis

Automatic thresholding in FIJI was used to identify cell areas in phase-contrast frames of time lapses, and non-cell areas were subtracted from frames. In time lapses of single cell growth, images were converted to binary images and objects were filtered according to a size minimum of 1000 px² (0.631 µm/px). In time lapses of cell growth within colonies, objects were filtered according to a size minimum of 5000 px². The 2D correlation coefficient is defined as 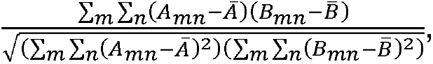 where *A* and *B* are *m* × *n* arrays with respective means A and B^B^.

Reading each frame as an array, the 2D correlation coefficient was calculated for 72 pairs of successive frames using the inbuilt function corr2 in MATLAB software (R2023a version, Mathworks).

### Epithelial features and Bayesian inference of epithelial forces

Bayesian force inference was calculated using previously published methodologies^40, 43^. Images of hiPSC epithelia marked with ZO1 were segmented using the Tissue Analyzer^63^. These segmentations were used to calculate each cell’s shape index, defined as 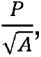 where P is the cell perimeter and A is the cell area^38^. Segmentations were exported into the Bayesian force inference MATLAB package (version R2023a) ^43^. To facilitate comparisons of forces between different images/conditions, images were stitched alongside each other in FIJI prior to segmentation and force inference, as in Figure 5B. Image size was chosen to balance maximizing data acquisition (≥135 cells per image), minimizing areas of disruption and minimizing high segmentation/computation time.

## Statistical analyses

All statistical analyses were performed using Prism software (8.4.3 version, GraphPad). Data are represented as mean ± SEM, and p values show the results of a two-tailed t test (*, < 0.05; **, < 0.01; ***, < 0.001). The number of replicates in *in vitro* measurements represents the number of gel-coated/glass coverslips tested. Where replicates were performed using cells of different passage number, replicates were paired accordingly. ZO1 organization experiments were found to be particularly sensitive to cells’ growth phase at the onset of experiments, and so replicates were ratio-paired. Statistical significance of cell motility was assessed by comparing 72 pairs of successive frames for each time lapse. Statistical significance of the shape index and of the results of Bayesian force inference was assessed by comparing the shape index, the inferred stress tensor magnitude along the axis of maximum stress, and the inferred pressure values of ≥135 cells in each image.

## Key resources table

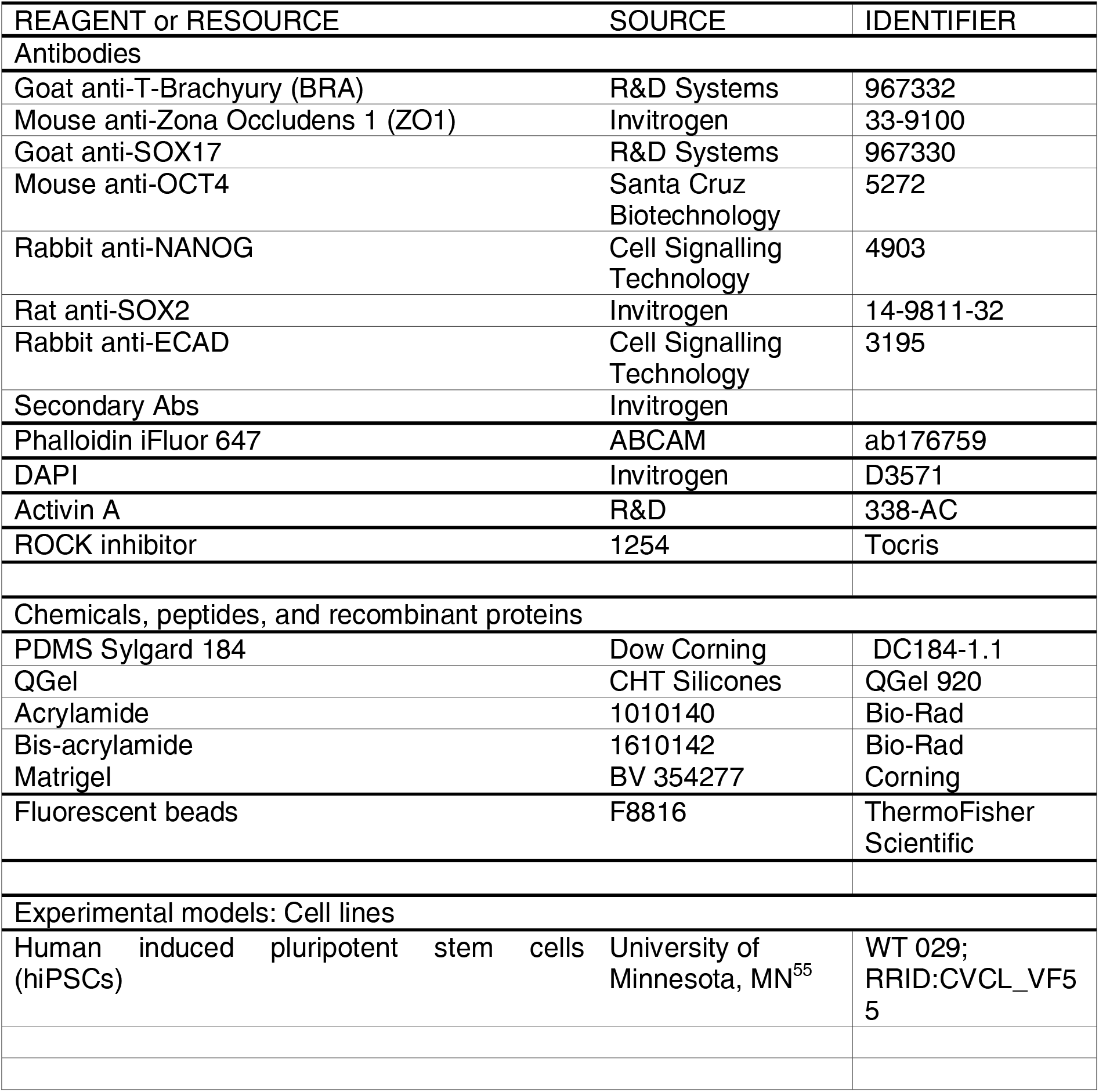
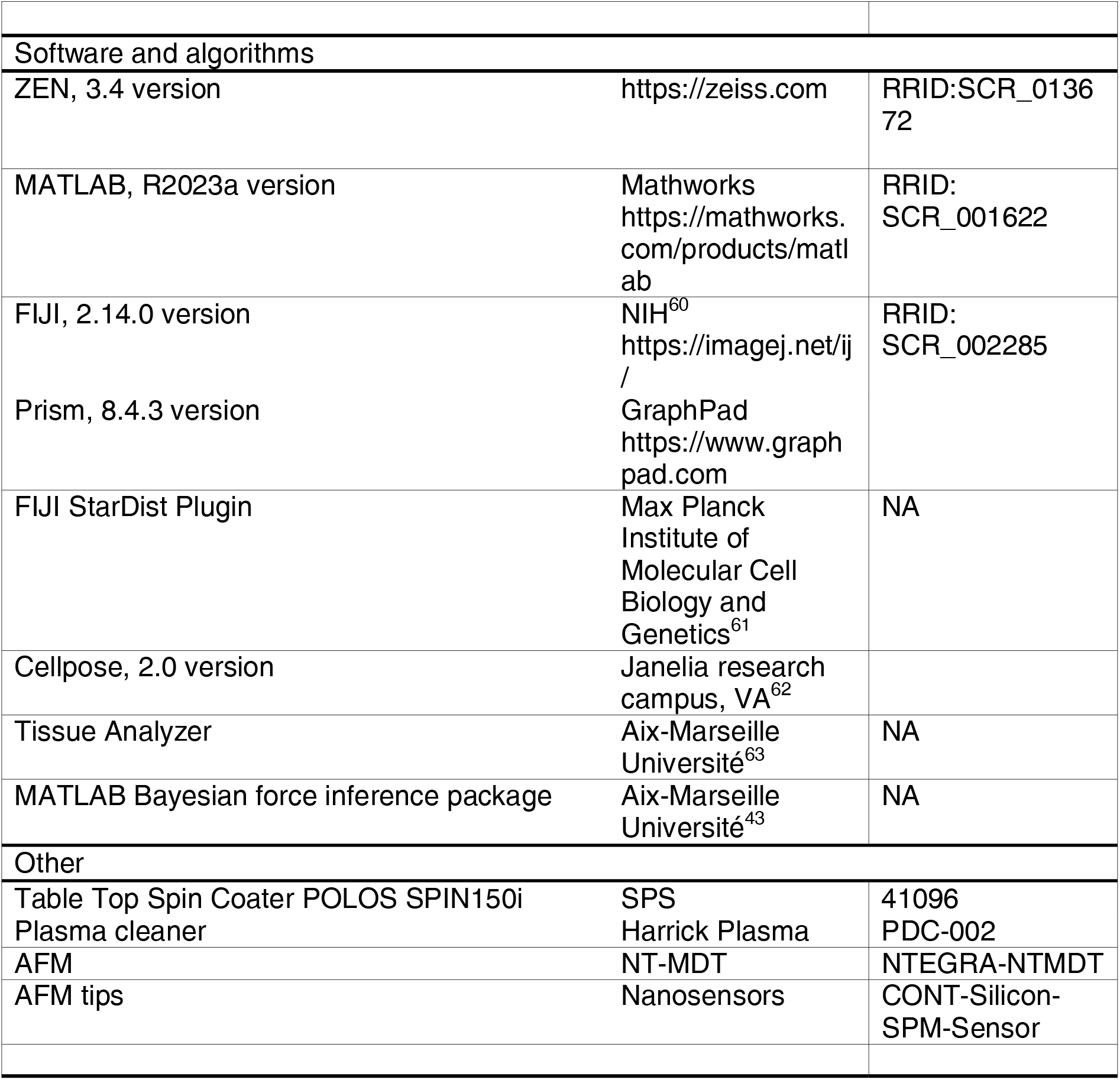

## Supporting information

Llewellyn et al 2024 Supplemental Information

Llewellyn et al 2024 Supplemental-Figure-1

Llewellyn et al 2024 Supplemental-Figure-2

Llewellyn et al 2024 Supplemental-Figure-3

## Acknowledgements

We thank: all members of our labs for helpful discussions and comments; L. Fasano and M. Merkel for critical reading of the manuscript; T. Legier and E. Bazellières for discussions and reagents, T. Legier and T. Boudier for image analysis pipelines and macros, T. Duong Quoc Le for assistance with video analysis, C. Guijarro-Calvo and W. Kong for assistance with Tissue Analyzer & Bayesian force inference algorithms. Microscopy was performed at the imaging platform of the IBDM, supported by the ANR through the “Investments for the Future” program (France- BioImaging, ANR-10-INSB-04-01). The authors thank members of the IBDM core facility for microscopy for their technical support. The IBDM is affiliated to NeuroMarseille, the AMU neuroscience network, and to NeuroSchool, the AMU graduate school in neuroscience supported by the A*MIDEX foundation (AMX-19-IET-004) and the “Investissements d’Avenir” program (nEURo*AMU, ANR-17-EURE-0029 grant). E.H. belongs to the French Consortium Approches Quantitatives du Vivant / Quantitative Approaches to Living Systems (AQV). J.L. was supported by the Turing Center for Living Systems (CENTURI), funded by France 2030, the French Government programme managed by the French National Research Agency (ANR-16-CONV-0001), and by the Excellence Initiative of Aix-Marseille University - A*MIDEX (A-M-AAP-ID-17-66-170301-11.30). This work was also funded by France Parkinson (Convention 2021-239736), Fondation de France (2023-265613), Fondation Louis Justin Besançon, COEN Pathfinder III (Network of Centres of Excellence in Neurodegeneration; COEN4014) to RD, and by Excellence Initiative of Aix-Marseille University - A*MIDEX (A-M-AAP-ID-17-66-170301-11.30) to EH. The funders had no role in study design, data collection and analysis, decision to publish, or preparation of the manuscript.

## Author contributions

J.L., E.H., and R.D. designed the research; J.L., A.C. conducted experiments; J.L., E.H., and R.D. analyzed data; J.L., A.C. produced initial images for figures; and J.L. and R.D. wrote the paper with assistance from all authors. J.L., A.C., E.H., and R.D. reviewed and edited the manuscript.

E.H. and R.D. provided funding.

## Supplemental Figures

**Figure S1.** Control of substrate stiffness through modifying gel composition preserves hiPSC pluripotency and facilitates hiPSC growth and differentiation, down to a lower stiffness limit.

**(A)** Representative IF images of the nuclear marker DAPI and of the pluripotency markers OCT4 and E-CAD in hiPSCs grown on glass (top) and PDMS reticulated using different concentrations of cross linker (5%, 4%, 3%, 2% from rows 2-5, respectively). Cells were fixed and images were taken 24 hours after the removal of ROCK inhibitor (48 hours after cell seeding at 110,000 cells/cm^2^).

**(B)** Phase contrast images of hiPSC colonies growing on glass (top) and 77.6 kPa PDMS gel (bottom) at time points 24 and 30 hours after the removal of ROCK inhibitor (48 and 54 hours after cell seeding at 15,000 cells/cm^2^). Taken from the first and final time points of Supplemental Movies 3 and 4.

**(C)** Confluency, defined as the percentage of the frame of view covered by cell colonies, of hiPSCs grown on glass (black) and 77.6 PDMS gel (purple) over time. Images were taken 5 min apart, 24-30 hours after the removal of ROCK inhibitor (48-54 hours after cell seeding at 15,000 cells/cm^2^) and a 30 min moving average is shown. Frames of view were selected for similar confluency values (32% and 33%) at the first time point.

**(D)** Brightfield images of the capability/incapability of hiPSCs to adhere and grow as a 2D epithelium on 77.6 kPa stiffness PDMS, 25.1 kPa PA, 11.6 kPa PA, and 4.07 kPa PA. Cells were fixed and images were taken 24 hours after cell seeding at 110 k cells/cm^2^.

**(E)** Representative IF images of DAPI and of the mesendoderm marker BRA in hiPSCs grown on 77.6 kPa PDMS gel (left), and 11.6 kPa polyacrylamide (PA) hydrogel (right). Cells were fixed and images were taken 24 hours after the addition of 100 ng/ml Activin A (48 hours after the removal of ROCK inhibitor, and 72 hours after cell seeding at 110,000 cells/cm^2^).

**(F)** Quantification of the number of BRA-positive cells in the three stiffness conditions at the time point illustrated in **(C)**. BRA-positive cells were defined as those whose mean fluorescence intensity was above the 95^th^ percentile of the mean BRA intensity in Activin A-negative control cells at the same time point and stiffness. At least 10 randomly selected frames of view were taken per condition, covering >2.23 mm² in total.

Scale bar in all images = 100 µm. Data are represented as mean ± SEM, and p values show the results of a two-tailed t test (* < 0.05, ** < 0.01, *** < 0.001)

**Figure S2.** A lag in epithelial organization on PDMS gel substrates occurs independently of gel surface chemistry, gel topography, or gel composition.

**(A)** Representative IF images of the nuclear marker DAPI and of the tight junction protein ZO1 in hiPSCs grown on glass (left), 2.18 MPa stiffness PDMS, middle), and 77.6 kPa stiffness PDMS (right). Cells were fixed and images were taken at three different time points: 24 (top), 48 (middle), and 72 hours (bottom) after the removal of ROCK inhibitor (48, 72, and 96 hours after cell seeding at 110,000 cells/cm^2^ respectively).

**(B)** Representative IF images of DAPI and ZO1 in hiPSCs grown on glass (left), 2.18 MPa stiffness PDMS (middle), and 77.6 kPa stiffness PDMS (right). The top row shows PDMS gels prepared as normal, while the bottom row shows PDMS gels which have been plasma-coated to ensure high hydrophilicity of the gel surface. Cells were fixed and images were taken 24 hours after the removal of ROCK inhibitor (48 hours after cell seeding at 110,000 cells/cm^2^).

**(C)** (Left, x-y plane) representative IF images demonstrating the method used to investigate the surface topography of gel substrates. (Right, x-z plane, 10 µm in total height) fluorescent beads on the gel surface allow mapping of the height of the gel across the surface.

**(D)** The ZO1 organization, defined as the percentage of the frame of view covered by ZO1- bounded cells, in hiPSCs grown on glass, 32.1 kPa stiffness PA, and 77.6 kPa stiffness PDMS. Areas of ZO1-bounded cells were detected automatically. Cells were fixed and images were taken 48 hours after the removal of ROCK inhibitor (72 hours after cell seeding at 60,000 cells/cm^2^). At least 3 randomly selected frames of view were taken per condition, covering >5.28 mm² in total. N = 4 unpaired replicates.

Scale bar in all images = 100 µm unless otherwise specified. Data are represented as mean ± SEM, and p values show the results of a two-tailed t test (* < 0.05, ** < 0.01, *** < 0.001).

**Figure S3.** AFM measurements of Young’s modulus of gel substrates across a wide range of stiffnesses.

**(A)** Examples of loading/unloading AFM curves on PDMS gels, PA hydrogels, and QGels. Note that PDMS and PA hydrogel showed very elastic behaviors in the range of moduli that were investigated with loading (grey) and unloading (colored) curves perfectly superimposed with each other. The QGels showed some viscoelastic behavior with the unloading curves below the loading curve.

**(B)** Examples of loading curves for the whole set of investigated gels. For each type of gel, a number N of measurements was performed ranging from 23 to 51, on a total of 3 to 5 different samples. The resulting average Young’s moduli vary from 2 MPa for the stiffest PDMS sample to less than 1.9 kPa for the softest QGel sample.

**(C)** Schematic representation of the tissue showing stiffnesses comparable to the gels used in this study^64^.

## Supplemental Videos

### Video S1

Phase contrast time lapse of human induced pluripotent stem cells growing at low confluency on glass. Images taken at 5 min interval span time points from 0 to 6 hours after the removal of ROCK inhibitor (24 to 30 hours after cell seeding at 5,000 cells/cm^2^).

### Video S2

Phase contrast time lapse of human induced pluripotent stem cells growing at low confluency on 77.6 kPa PDMS gel. Images taken at 5 min interval span time points from 0 to 6 hours after the removal of ROCK inhibitor (24 to 30 hours after cell seeding at 5,000 cells/cm^2^).

### Video S3

Phase contrast time lapse of human induced pluripotent stem cells growing at low confluency on glass. Images taken at 5 min interval span time points from 24 to 30 hours after the removal of ROCK inhibitor (48 to 54 hours after cell seeding at 15,000 cells/cm^2^).

### Video S4

Phase contrast time lapse of human induced pluripotent stem cells growing at low confluency on 77.6 kPa PDMS gel. Images taken at 5 min interval span time points from 24 to 30 hours after the removal of ROCK inhibitor (48 to 54 hours after cell seeding at 15,000 cells/cm^2^).

