## Supplementary figures and images for "Substrate Stiffness Reshapes Layer Architecture and Biophysical Features of Human Induced Pluripotent Stem Cells to Modulate their Differentiation Potential"

### Llewellyn et al 2024 Supplemental-Figure-1

#### Supplemental Figure 1

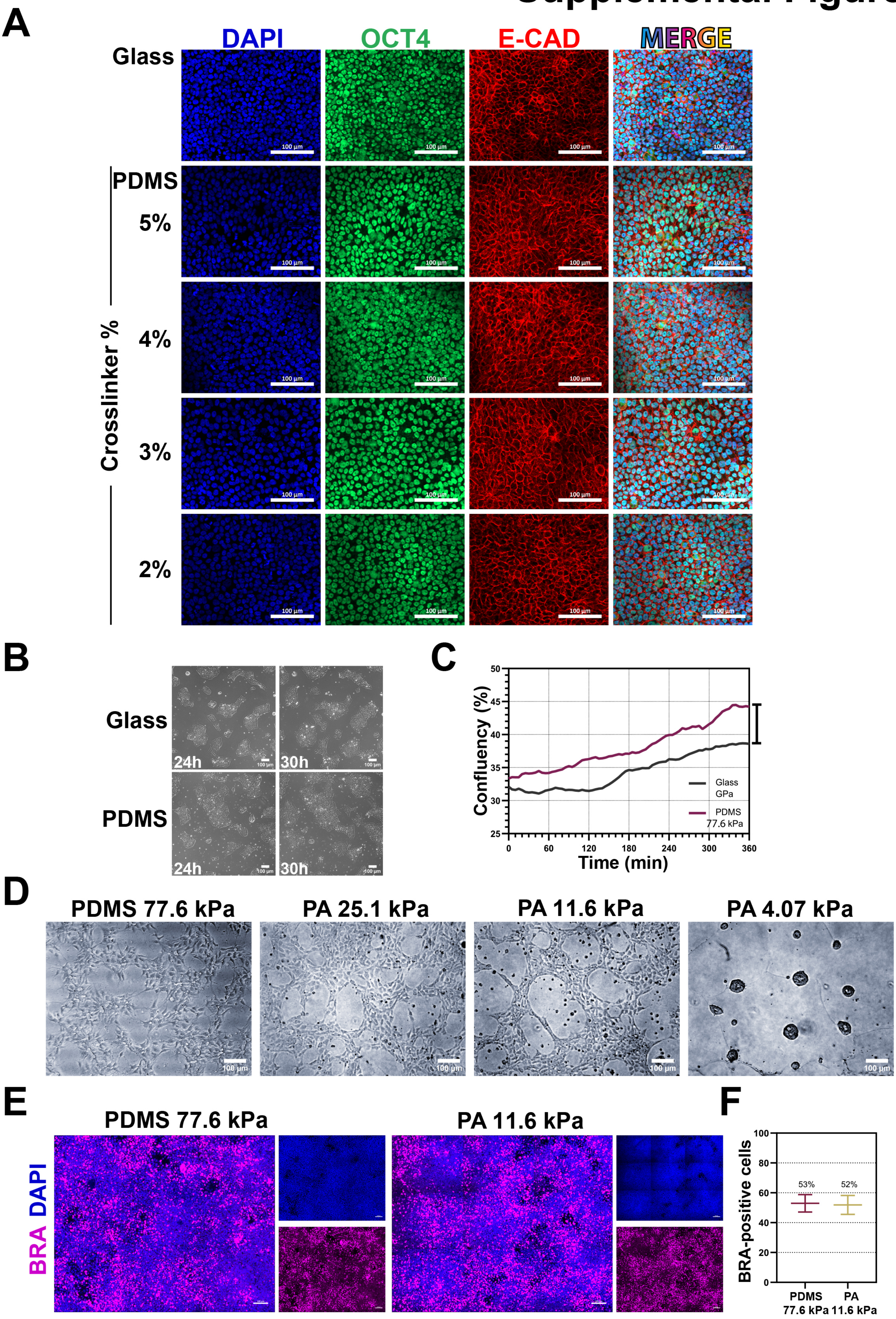

### Llewellyn et al 2024 Supplemental-Figure-2

#### Supplemental Figure 2

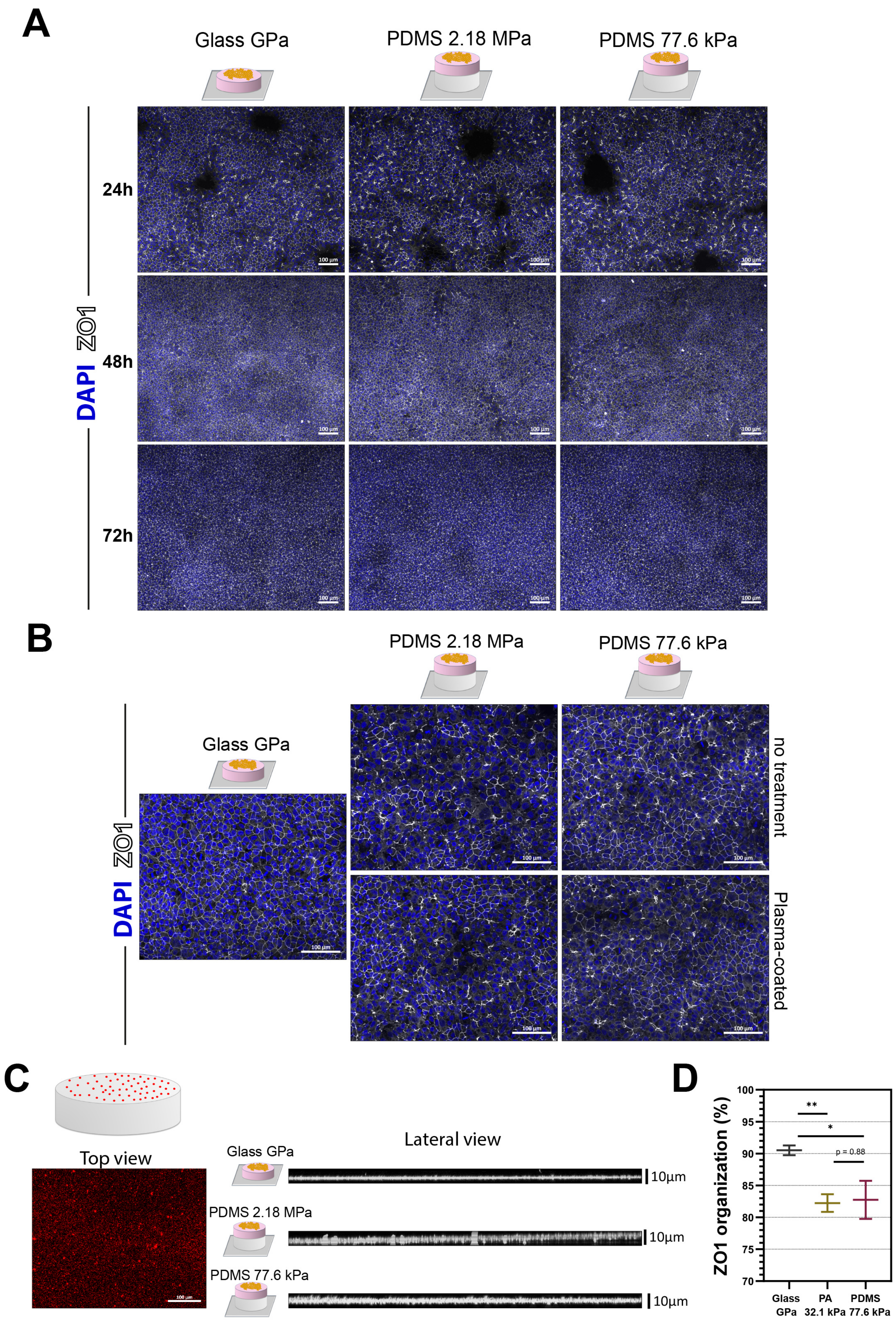

### Llewellyn et al 2024 Supplemental-Figure-3

#### Supplemental Figure 3

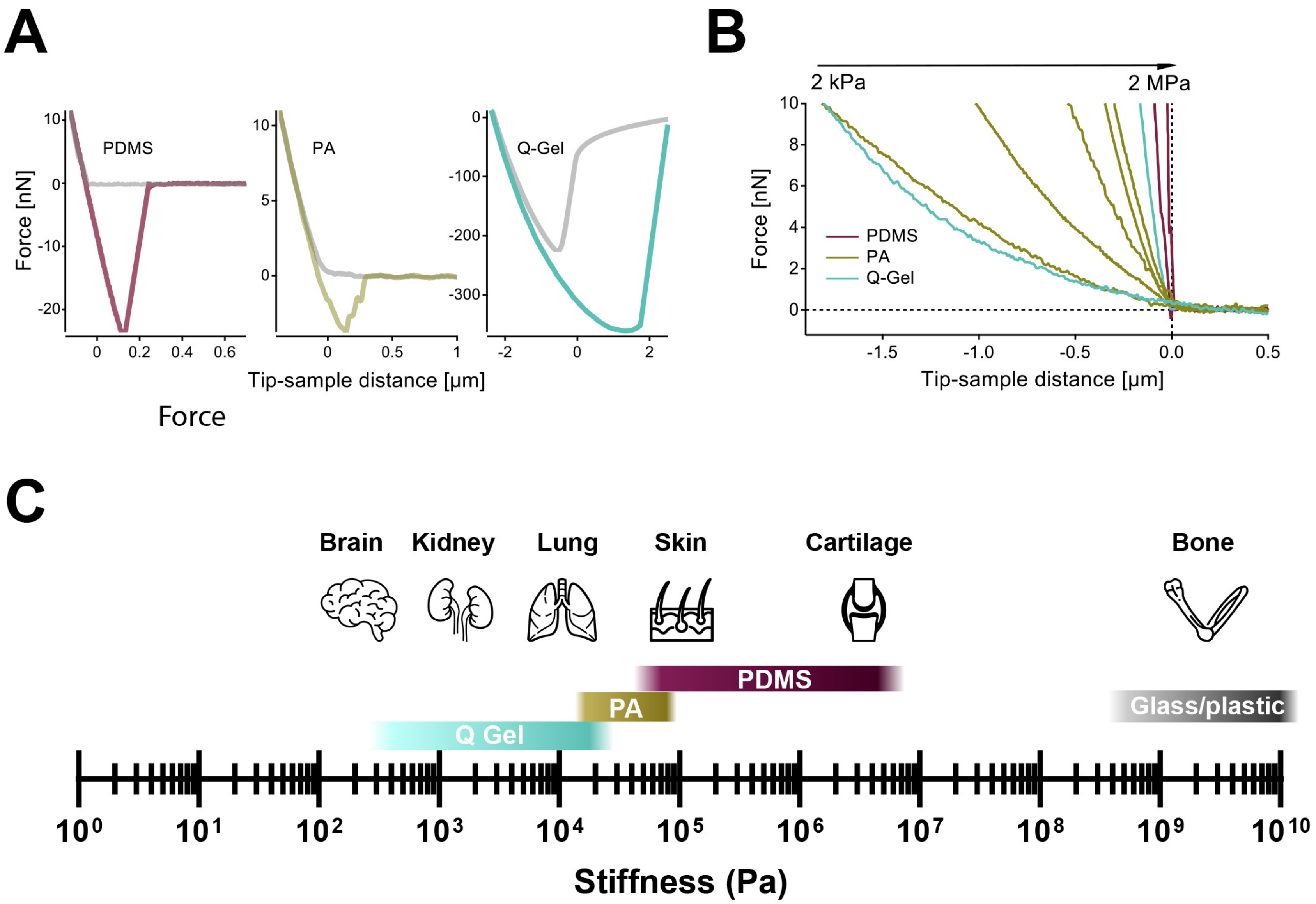
